# A compendium of genetic regulatory effects across pig tissues

**DOI:** 10.1101/2022.11.11.516073

**Authors:** The FarmGTEx-PigGTEx Consortium, Jinyan Teng, Yahui Gao, Hongwei Yin, Zhonghao Bai, Shuli Liu, Haonan Zeng, Lijing Bai, Zexi Cai, Bingru Zhao, Xiujin Li, Zhiting Xu, Qing Lin, Zhangyuan Pan, Wenjing Yang, Xiaoshan Yu, Dailu Guan, Yali Hou, Brittney N. Keel, Gary A. Rohrer, Amanda K. Lindholm-Perry, William T. Oliver, Maria Ballester, Daniel Crespo-Piazuelo, Raquel Quintanilla, Oriol Canela-Xandri, Konrad Rawlik, Charley Xia, Yuelin Yao, Qianyi Zhao, Wenye Yao, Liu Yang, Houcheng Li, Huicong Zhang, Wang Liao, Tianshuo Chen, Peter Karlskov-Mortensen, Merete Fredholm, Marcel Amills, Alex Clop, Elisabetta Giuffra, Jun Wu, Xiaodian Cai, Shuqi Diao, Xiangchun Pan, Chen Wei, Jinghui Li, Hao Cheng, Sheng Wang, Guosheng Su, Goutam Sahana, Mogens Sandø Lund, Jack C.M. Dekkers, Luke Kramer, Christopher K. Tuggle, Ryan Corbett, Martien A.M. Groenen, Ole Madsen, Marta Gòdia, Dominique Rocha, Mathieu Charles, Cong-jun Li, Hubert Pausch, Xiaoxiang Hu, Laurent Frantz, Yonglun Luo, Lin Lin, Zhongyin Zhou, Zhe Zhang, Zitao Chen, Leilei Cui, Ruidong Xiang, Xia Shen, Pinghua Li, Ruihua Huang, Guoqing Tang, Mingzhou Li, Yunxiang Zhao, Guoqiang Yi, Zhonglin Tang, Jicai Jiang, Fuping Zhao, Xiaolong Yuan, Xiaohong Liu, Yaosheng Chen, Xuewen Xu, Shuhong Zhao, Pengju Zhao, Chris Haley, Huaijun Zhou, Qishan Wang, Yuchun Pan, Xiangdong Ding, Li Ma, Jiaqi Li, Pau Navarro, Qin Zhang, Bingjie Li, Albert Tenesa, Kui Li, George E. Liu, Zhe Zhang, Lingzhao Fang

## Abstract

The Farm animal Genotype-Tissue Expression (FarmGTEx, https://www.farmgtex.org/) project has been established to develop a comprehensive public resource of genetic regulatory variants in domestic animal species, which is essential for linking genetic polymorphisms to variation in phenotypes, helping fundamental biology discovery and exploitation in animal breeding and human biomedicine. Here we present results from the pilot phase of PigGTEx (http://piggtex.farmgtex.org/), where we processed 9,530 RNA-sequencing and 1,602 whole-genome sequencing samples from pigs. We build a pig genotype imputation panel, characterize the transcriptional landscape across over 100 tissues, and associate millions of genetic variants with five types of transcriptomic phenotypes in 34 tissues. We study interactions between genotype and breed/cell type, evaluate tissue specificity of regulatory effects, and elucidate the molecular mechanisms of their action using multi-omics data. Leveraging this resource, we decipher regulatory mechanisms underlying about 80% of the genetic associations for 207 pig complex phenotypes, and demonstrate the similarity of pigs to humans in gene expression and the genetic regulation behind complex phenotypes, corroborating the importance of pigs as a human biomedical model.

## Introduction

Genome-wide association studies (GWAS) and other technological advances are revealing genomic variants associated with complex traits and adaptive evolution at an unprecedented speed and scale in both plants^1^ and animals^2^, but particularly in humans^3,4^. However, most of the variants fall in non-coding regions, putatively contributing to phenotypic variation by regulating gene activity/structure at different biological levels (e.g., tissues and cell types)^5,6^. The systematic characterization of genetic regulatory effects on transcriptome (e.g., expression quantitative trait loci, eQTL) across tissues, as carried out in the Genotype-Tissue Expression (GTEx) project in humans^7^, has proven to be a powerful strategy for connecting GWAS loci to gene regulatory mechanisms at large scale^8–10^.

To sustain food and agriculture production while minimize associated negative environmental impacts, it is crucial to identify molecular mechanisms that underpin complex traits of economic importance in farm animals to enable biology-driven breeding biotechnologies. However, the annotation of regulatory variants in farm animals has so far been limited by sample size, tissue/cell type, and genetic background^11–13^. We therefore launched the international Farm animal GTEx (FarmGTEx, https://www.farmgtex.org/) project to build a comprehensive open-access atlas of regulatory variants in domestic animal species. This resource along with the Functional Annotation of Animal Genomes project (FAANG) will not only facilitate fundamental biology discovery but also enhance the genetic improvement of farm animal species^14^.

Pigs are an important agricultural species by supplying animal protein resource for humans, and serve as an essential biomedical model for studying human development, disease, organ xenotransplantation, and drug design due to their similarity to humans in multiple attributes such as genome, anatomical structure, physiology and immunology^15^. Here we report the first results of the PigGTEx which is underpinned by 9,530 RNA-Seq data from 138 tissue/cell types and 1,602 whole-genome sequence (WGS) samples from over 100 pig breeds collected around the world. We test the association of transcriptomic phenotypes with 3,087,268 DNA variants in 34 pig tissues with sufficient sample sizes (n > 40), and then evaluate the tissue- and breed-sharing patterns of detected regulatory effects. We examine sequence ontology, epigenetic modifications, and chromatin conformation to identify putative molecular mechanisms underlying regulatory impacts of the associated variants. We then apply this resource to functionally dissect GWAS associations for 268 complex traits through multiple complementary integrative methods such as transcriptome-wide association studies (TWAS), colocalization, and Mendelian randomization (MR). Finally, we leverage the human GTEx resource and GWAS summary statistics of 136 human complex traits and diseases to assess the similarity between pigs and humans in both gene expression and regulation, and elucidate their impacts on complex phenotypes. We make the PigGTEx resources freely accessible and easy-to-use *via* http://piggtex.farmgtex.org. PigGTEx resources will be valuable tools to increase our understanding of complex trait genetics, domestication, and evolutionary adaptation in pig, enhance breeding biotechnology and veterinary medicine, and benefit human biomedical research.

## Results

### Data summary

We uniformly processed 9,530 RNA-Seq samples, yielding a total of ~280 billion clean reads with an average mapping rate of 80.33%. Details of these samples are summarized in Fig. S1 and Table S1. After filtering out samples of low quality (see Methods), we retained 7,095 RNA-Seq samples for downstream analysis. We quantified expression levels for protein coding genes (PCG), lncRNA, exons and enhancers, and alternative splicing events in these samples. Sample clustering based on the five transcriptomic phenotypes recapitulated tissue types well, implying that pig public RNA-Seq data can reflect the tissue-specific biology after processing uniformly (Fig. 1a-b and Fig. S2), concordant with our previous findings in cattle^16^.

**Figure 1.**
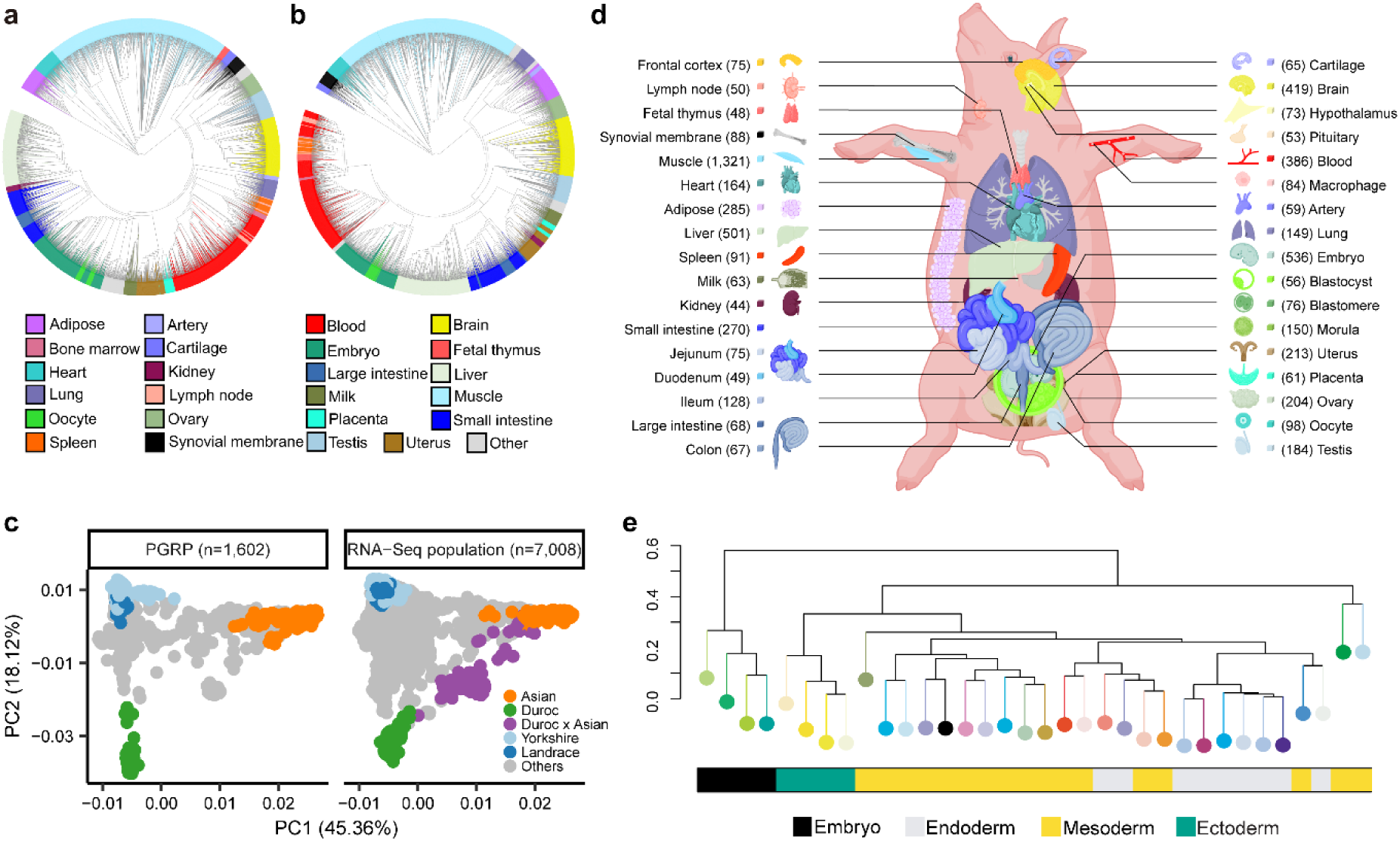
Characteristics of samples in the pilot phase of PigGTEx project. (**a**) Phylogenetic clustering of 7,095 RNA-Seq samples based on the normalized expression (Transcripts Per Million, TPM) of 6,500 highly variable genes, defined as the top 20% of genes with the largest standard deviation of TPM across samples. (**b**) The same sample clustering as (**a**) but based on normalized alternative splicing values (Percent Spliced In, PSI) of 6,500 highly variable spliced introns, defined as the top 13% of spliced introns with the largest standard deviation of PSI across samples. (**c**) Principal component analysis of samples based on 12,207 LD-independent (*r*^2^ < 0.2) SNPs. The left panel is for whole-genome sequencing samples (n = 1,602) in the Pig Genomics Reference Panel (PGRP), while the right one is for RNA-Seq samples (n = 7,008) with successful genotype imputations. (**d**) Sample sizes of 34 tissues, cell types and organ systems (all referred to as “tissues”) used for molecular quantitative trait loci mapping. (**e**) Hierarchical clustering of 34 tissues based on the median expression of all 31,871 Ensembl annotated genes (v100) across samples within tissues, representing embryo, endodermal, mesodermal, and ectodermal lineages.

We called a median number of 74,347 SNPs directly from the 7,095 RNA-Seq samples (Extended Data Fig. 1a-b). Leveraging a newly built large multi-breed Pig Genomics Reference Panel (PGRP) consisting of 1,602 WGS samples (1,093 publicly available and 509 newly generated) (Fig. S3 and Table S2), we imputed genotypes of RNA-Seq samples to the WGS level with an imputation accuracy of over 95% across multiple validation strategies (Extended Data Fig. 1c-h). We also predicted missing breed information for 3,684 (52.6%) public RNA-Seq samples using their imputed genotypes (Extended Data Fig. 1i-m, and Table S1). Sample clustering based on genotypes overlapping both datasets demonstrated that the population structure of the RNA-Seq samples was similar to the PGRP (Fig. 1c). After removing duplicated RNA-Seq samples based on their genetic relatedness (identity by state, IBS > 0.9) derived from imputed genotypes, we retained a total of 5,457 samples representing 34 tissues, cell types, or organ systems (all referred to as ‘tissues’ hereafter), with at least 40 samples per tissue, for subsequent analysis (Fig. 1d, Extended Data Fig. 2a-e). The 34 tissues originated from embryos and all three germ layers, and represent all major lineages in the pig body (Fig. 1e). The number of detected molecular phenotypes in each of 34 tissues are summarized in Table S3. We further analyzed 270 public pig multi-omics datasets, including 245 whole genome bisulfite sequence (WGBS) (Fig. S4-5 and Table S4-6), 20 single-cell RNA-seq (Fig. S6 and Table S7) and five Hi-C samples (Table S8-9), for the following integrative analysis.

### The gene expression atlas empowers functional annotation of pig genes

Gene expression was either tissue-specific or ubiquitous (Fig. S7a). However, compared to non-coding genes, the expression of PCG was less tissue-specific (Extended Data Fig. 3a). We detected between 145 (morula) and 5,180 (frontal cortex) tissue-specific expressed genes acorss 34 tissues (Extended Data Fig. 3b, Fig. S7b). The functions of tissue-specific genes clearly reflected the known biology of their respective tissues (Extended Data Fig. 3b). By examining previously predicted chromatin states^17^(Fig. S7c), we found that tissue-specific genes showed a higher enrichment of active regulatory elements and a higher depletion of repressed polycomb regions in matching tissues than in nonmatching tissues (Extended Data Fig. 3c-e). For instance, *MYL2*, that plays a key role in heart development and function, showed heart/muscle-specific gene expression and promoter activity (Extended Data Fig. 3d), as did *MYOG* in muscle and *APOH* in liver (Fig. S7d). In addition, tissue-specific genes exhibited distinct patterns of evolutionary DNA sequence constraints across tissues (Fig. S7e), in agreement with the hypothesis of tissue-driven evolution^18^.

To assign function to pig genes, we employed the principle of ‘guilt by association’ by performing a gene co-expression analysis in each of the 34 tissues (Fig. S8a). In total, we detected 5,309 co-expression modules across tissues, and assigned 25,023 genes to at least one module (Fig. S8b-c, Table S10). Among them, 13,266 (42.57%) genes had no functional annotation in the Gene Ontology (GO) database (Extended Data Fig. 3f, Fig. S8d); these are referred to as “unannotated genes” hereafter. Unannotated genes were less expressed, showed weaker DNA sequence conservation, lower proportion of orthologous genes, and higher tissue-specificity than genes with functional annotations (Extended Data Fig. 3f). The proportion of expressed unannotated genes varied across tissues, indicating differences in functional annotation between tissues (Extended Data Fig. 3g). The frontal cortex and hypothalamus had the largest proportion of unannotated genes (an average of 29.51% and 28.91%, respectively), whereas morula and embryo had the smallest (an average of 10.10% and 13.24%, respectively). For instance, 42 unannotated genes were co-expressed with 59 functional annotated genes in pituitary, which were significantly enriched in neuron apoptotic processes (Extended Data Fig. 3h). Altogether, these results suggest that the PigGTEx gene expression atlas serves as a useful resource to enhance the functional annotation of the pig reference genome.

### Molecular QTL (molQTL) mapping

To explore the genetic determinants underlying variation in gene expression, we first estimated the *cis*-heritability (*cis*-*h^2^*) for PCG expression while accounting for estimated confounding factors (Extended Data Fig. 2f-h), yielding an average *cis*-*h*^2^ of 0.14 across the 34 tissues (Extended Data Fig. 4a). Tissues with similar biological functions clustered together based on the *cis*-*h^2^* estimates (Extended Data Fig. 4b). We then mapped molQTL for five molecular phenotypes in each tissue, including *cis*-eQTL for PCG expression, *cis*-eeQTL for exon expression, *cis*-lncQTL for lncRNA expression, *cis*-enQTL for enhancer expression, and *cis*-sQTL for alternative splicing. We found that 86%, 67%, 46%, 27% and 52% of all tested PCGs (n = 17,431), lncRNAs (n = 7,374), exons (n = 82,678), enhancers (n = 128,600) and genes with alternative splicing events (n = 18,331) had at least one significant variant (eVariant) (FDR < 0.05) detected in at least one tissue; hence they were defined as eGenes, eLncRNAs, eExons, eEnhancers and sGenes, respectively (Table S11). The proportion of eGenes detected was strongly and positively correlated (Pearson’s *r* = 0.81, *P* = 6.86×10^−9^) with sample size across tissues (Fig. 2a). A similar pattern was observed for the other four molecular phenotypes (Fig. S9). The top *cis*-e/sQTL centered around transcription start sites (TSS) of genes (Fig. S10a-g). Tissues with a larger sample size yielded a larger proportion of *cis*-eQTL with small effects (Fig. S10h-i). In general, PCG had the highest proportion of detected eGenes across tissues, followed by lncRNA, splicing, exon and finally enhancer (Fig. 2b). Notably, molecular phenotypes exhibited a high proportion (an average of 70%) of their own specific molQTL after taking linkage disequilibrium (LD) between SNPs into account (Fig. 2b), indicative of their distinct underlying genetic regulation.

**Figure 2.**
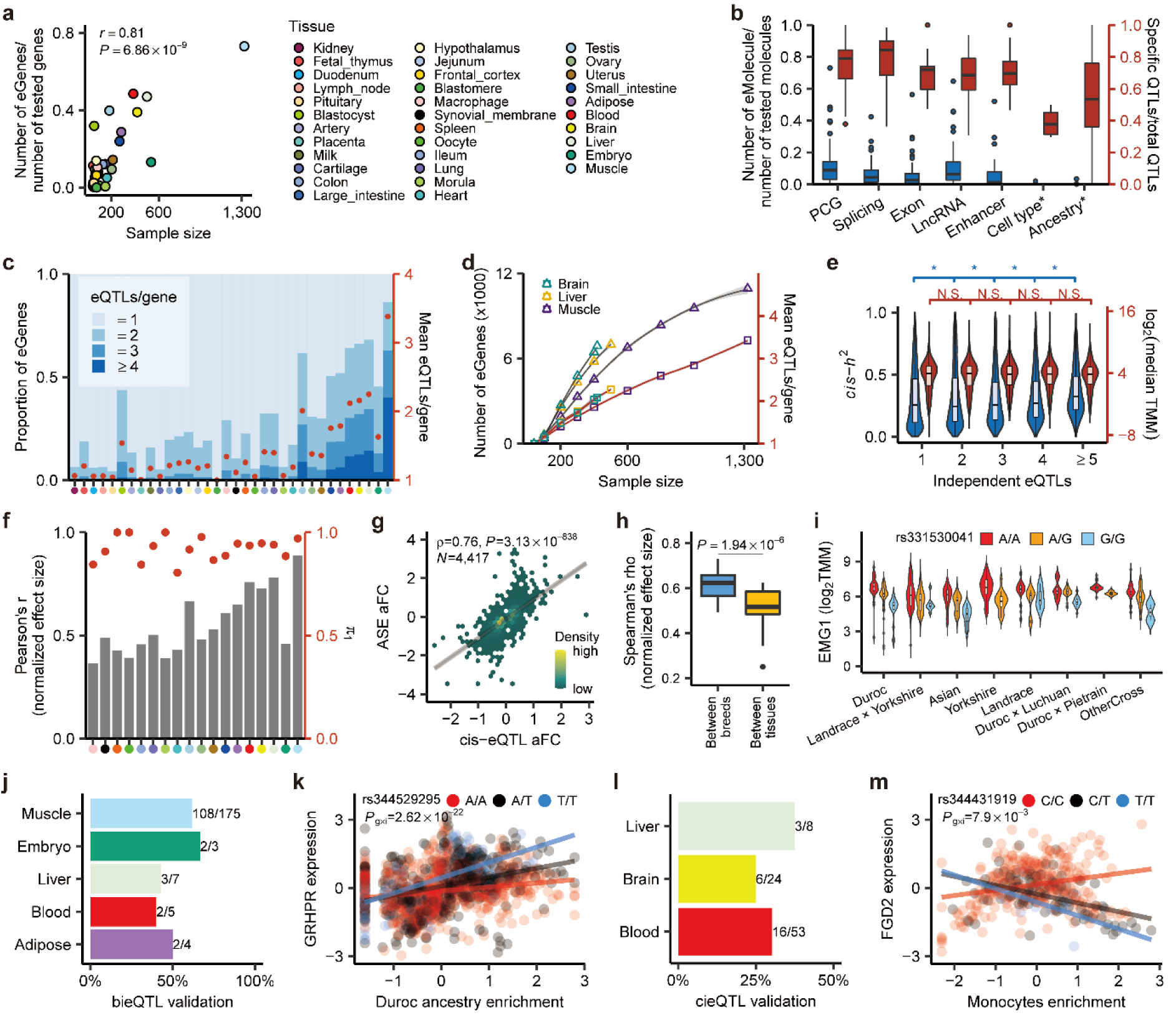
Molecular quantitative trait loci (molQTL) discovery. (**a**) Pearson’s correlation (*r*) between the proportion of detectable eGenes and sample size across 34 tissues. (**b**) Proportions of detectable eMolecule (blue, left) and specific molQTL (red, right) for seven types of molecular phenotypes in 34 tissues. Cell type* and Ancestry* are for cell-type and breed interaction QTLs (cieQTL and bieQTL), respectively. (**c**) Distribution and average number of independent *cis*-eQTL per gene across 34 tissues. Tissues (*x*-axis) are ordered (from smallest to largest) by sample size. The color key is the same as in (**a**). (**d**) Number of eGenes (left, triangle) and average number of independent *cis*-eQTL (right, square) in the down-sampling analysis. The smoothed lines are fitted by a local polynomial regression model. (**e**) The comparison of *cis*-heritability (*cis*-*h*^2^) (blue, left) and median expression levels (red, right) of genes with different numbers of detectable independent *cis*-eQTL across 34 tissues. TMM: Trimmed Mean of M-value normalized expression levels. * indicates *P* < 0.05 and N.S. indicates *P* > 0.05, based on the one-sided Student’s *t*-test. (**f**) Internal validation of *cis*-eQTL. Bars represent the Pearson’s *r* of normalized effect size of *cis*-eQTL between validation and discovery groups. Points represent the π_1_ statistic measuring the replication rate of *cis*-eQTL. (**g**) Spearman’s *ρ* of effect sizes (allelic fold change, aFC) between *cis*-eQTL and allele specific expression (ASE) at matched loci (n = 4,417) in muscle. (**h**) Comparison of Spearman’s *ρ* of normalized effect size of *cis*-eQTL between breeds with those between tissues. The *P* value is obtained from two-sided Student’s *t*-test. (**i**) Expression levels of *EMG1* gene in muscle across three genotypes of eVariant (rs331530041) in eight breed groups. (**j**) Proportion of bieQTL that are validated with the ASE approach. The numbers besides the bars represent numbers of validated bieQTL (left) and total bieQTL being tested (right). (**k**) Effect of eVariant (rs344529295) of *GRHPR* significantly interacted with the Duroc ancestry enrichment in muscle. (**l**) Proportion of cieQTL that are validated by the ASE approach. (**m**) Effect of eVariant (rs344431919) of *FGD2* significantly interacted with monocytes enrichment in blood. The lines are fitted by a linear regression model.

We further conducted conditionally independent molQTL mapping. On average, 20% of eGenes, 13.5% of sGenes, 21.2% of eExons, 23.5% of eLncRNAs, and 17% of eEnhancers had more than one independent eVariant across tissues, and the proportion increased with an increasing sample size of tissues (Fig. 2c, Extended Data Fig. 5a-d). Down-sampling analysis in the three major tissues (i.e., muscle, liver, and brain) further confirmed an impact of sample size on the statistical power for *cis*-eQTL discovery (Fig. 2d). Approximately half of the independent *cis*-eQTL were located within ±182Kb of TSS, and those with larger effect size were closer to TSS (Extended Data Fig. 5e-g). The eGenes with more independent *cis*-eQTL have a higher *cis*-*h^2^*, but no significant differences for the median gene expression level (Fig. 2e).

We applied four distinct strategies to validate the *cis*-eQTL. First, we observed that the summary statistics of *cis*-eQTL derived from the linear regression model in TensorQTL^19^ had a strong correlation (an average Pearson’s *r* of 0.91 across tissues) with those from a linear mixed model (Extended Data Fig. 6a). Second, we performed an internal validation in 18 tissues with over 80 samples by randomly dividing samples into two equal groups, and then conducting *cis*-eQTL mapping separately in both subgroups. This approach yielded a high replication rate (measured by π_1_) for *cis*-eQTL, with an average π_1_ of 0.92 (range 0.80 – 1.00) and an average of 0.56 (range 0.36 – 0.89) for Pearson’s *r* between effect sizes across tissues (Fig. 2f). Third, we found that 92%, 74%, 73%, and 69% of *cis*-eQTL in blood, liver, duodenum, and muscle, respectively, were replicated in independent datasets, but the replication rates depended on the sample size of validation populations (Extended Data Fig. 6b-d). Fourth, we further found that effects (allelic fold changes, aFC) derived from allele specific expression (ASE) analysis were significantly correlated with those from *cis*-eQTL mapping consistently across tissues (Extended Data Fig. 6e-g). For instance, in muscle, ASE-derived effects of 4,417 SNPs were significantly correlated (Spearman’s *ρ* = 0.76, *P* < 1×10^−300^) with their *cis*-eQTL effects (Fig. 2g). All these results demonstrated the high replicability of *cis*-eQTL in the established PigGTEx resource.

### Breed-sharing patterns of cis-eQTL

To understand how *cis*-eQTL are shared across pig breeds, we considered muscle as an example. We divided all 1,302 muscle samples into eight breed groups (all referred to as ‘breeds’ hereafter) and performed *cis*-eQTL mapping separately (Extended Data Fig. 7a, Table S12). Across all eight breeds, we detected 9,548 unique *cis*-eGenes, of which 97.1% could be replicated (M-value > 0.95, METASOFT^20^) in at least two of these breeds. For instance, *EMG1*, *NMNAT1* and *COMMD10* showed consistent *cis*-eQTL in muscle across all eight breeds (Fig. 2i, Extended Data Fig. 7b-c). The replication rates were higher in breeds with more samples (Extended Data Fig. 7d). For instance, the Landrace × Yorkshire crossbreed had the largest sample size (n = 374) replicated on average 95.6% of the *cis*-eQTL detected in the other seven breeds, whereas Landrace had the smallest sample size (n = 49) and consequently replicated only approximately half of *cis*-eQTL detected in the other breeds (Extended Data Fig. 7d). The *cis*-eQTL effects were positively correlated between breeds as well as clearly separated from other tissues (Fig. 2h, Extended Data Fig. 7e). In addition, effects of *cis*-eQTL from the multi-breed meta-analysis were strongly and significantly correlated (Pearson’s *r* = 0.92, *P* < 1×10^−300^) with those from the combined muscle population (Extended Data Fig. 7f). Compared to the single-breed, the combined population detected 86.2% more *cis*-eQTL (Extended Data Fig. 7g). All these findings demonstrated that the majority of *cis*-eQTL are shared across breeds, and thus combining samples from different breeds might increase the discovery power of *cis*-eQTL and narrow down the number of putative causal variants by reducing LD among SNPs.

To explore how breed interacts with genotype to modulate gene expression, we conducted breed-interaction *cis*-eQTL (bieQTL) mapping. In total, we identified 589 genes with at least one significant bieQTL in 13 tissues (Fig. 2j-k, Table S13). For instance, in muscle, bieQTL of *GRHPR* and *CA14* had a significant interaction with the proportion of Duroc ancestry (Fig. 2k, Extended Data Fig. 7h), while bieQTL of *ATE1* had a significant interaction with the proportion of Yorkshire ancestry (Extended Data Fig. 7i). Furthermore, we conducted a cell-type deconvolution analysis in seven tissues, demonstrating the variation of cell type composition across bulk tissue samples (Extended Data Fig. 8a-g). We identified a total of 376 genes with at least one significant cell-type interaction *cis*-eQTL (cieQTL) in three tissues (Fig. 2l, Table S13). For instance, the cieQTL of *FGD2*, *SCRN2* and *HIBADH* significantly interacted with the enrichment of monocytes, CD2^−^ T cells and CD4^+^ T cells in blood, respectively (Fig. 2m, Extended Data Fig. 8h-i). In addition, we validated half of these bieQTL and cieQTL with the ASE approach^21^ (Fig. 2j,l, and Extended Data Fig. 8j-m).

### Tissue-sharing patterns of molQTL

To understand the relationship between the transcriptome and its genetic regulation between tissues, we quantified the pairwise similarity of the 34 tissues based on 10 data types, including the five types of molQTL and the respective molecular phenotypes (Fig. 3a, Extended Data Fig. 9a-d). As observed for *cis*-*h^2^*, tissues with similar functions clustered together, and the tissue relationship was consistent across all 10 data types (Fig. 3b). This implied that tissues with similar transcriptional profiles are potentially driven by shared genetic regulatory effects, indicative of the shared underlying biological processes. Embryonic tissues, oocyte and testis formed a separate cluster, while the remaining tissues demonstrated a relatively high degree of similarity to one another (Fig. 3a). Among these tissues, uterus and small intestine showed the highest degree of similarity to other tissues, indicating that they might serve as a proxy for many tissues to study common genetic regulation (Fig. 3a). However, these tissues cannot be obtained without invasive procedures. The most easily accessible samples, i.e., blood and milk cells, showed an average correlation of 0.51 *cis*-eQTL effects with other tissues. Both had the highest similarity to immune tissues, followed by intestinal tissues, and finally testis and embryonic tissues. The overall tissue sharing of molQTL showed a U-shaped curve (Fig. 3c), demonstrating that the genetic regulation of the transcriptome tends to be either highly tissue-specific or ubiquitous. Among these molQTL, *cis*-eQTL of PCG had the highest degree of tissue sharing, followed by *cis*-lncQTL, *cis*-sQTL, *cis*-eeQTL and finally *cis*-enQTL (Fig. 3c, Extended Data Fig. 9e).

**Figure 3.**
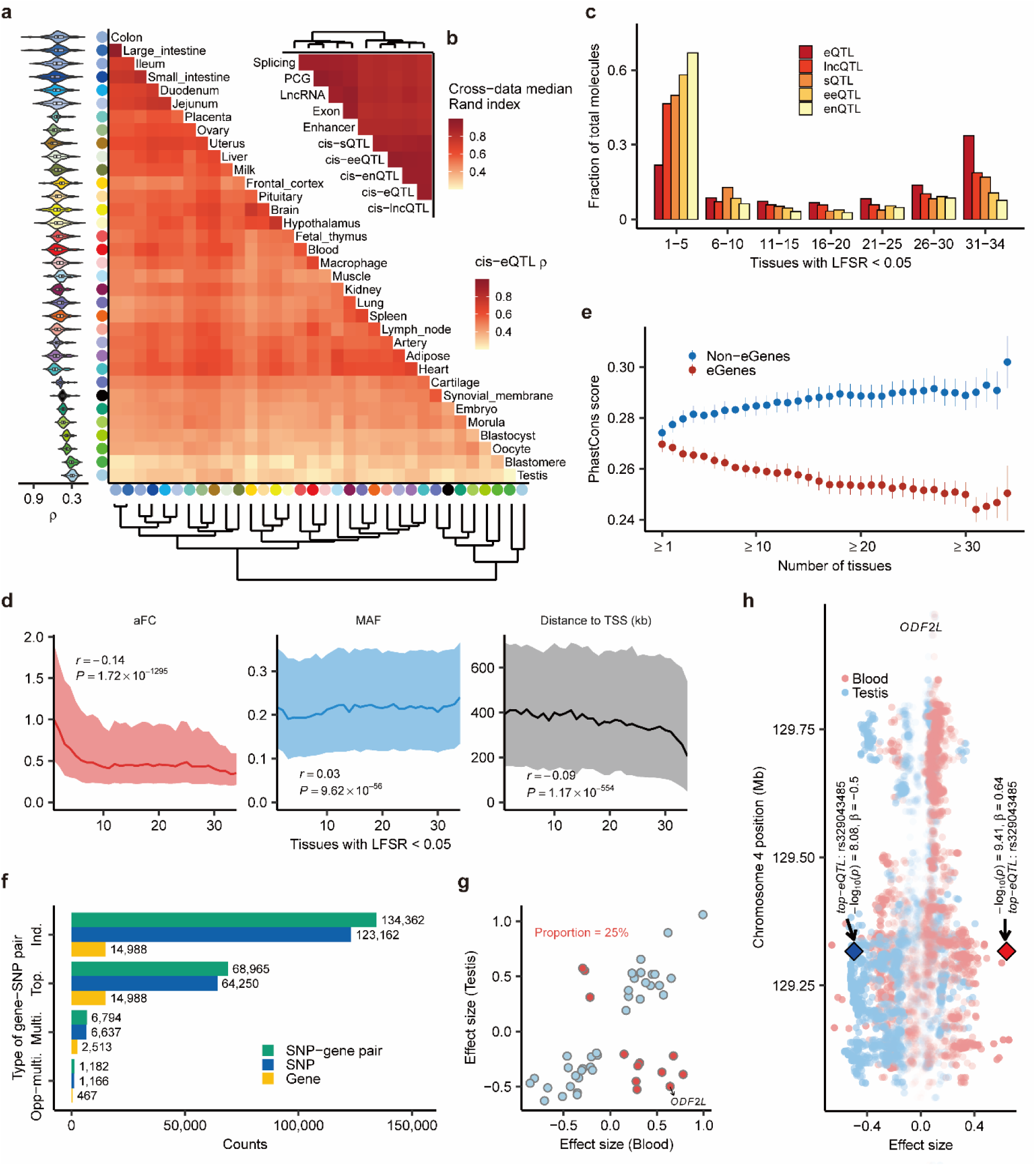
Tissue-sharing pattern of regulatory effects. (**a**) Heatmap of tissues depicting the corresponding pairwise Spearman’s correlation (*ρ*) of *cis*-eQTL effect sizes. Tissues are grouped by the hierarchical clustering (bottom). Violin plots (left) represent the Spearman’s *ρ* between the target tissue and other tissues. (**b**) Similarity (measured by the median pairwise Rand index) of tissue-clustering patterns across 10 data types. (**c**) The overall tissue-sharing pattern of five molQTL types at local false sign rate (LFSR) < 5% obtained by MashR (v0.2-6). (**d**) Relationships between the magnitude of tissue-sharing of *cis*-eQTL and their effect sizes (allelic fold change, aFC, left), minor allele frequencies (MAF, middle), and distances to transcript start site (TSS, right). The *P*-values are obtained by the Pearson’s correlation (*r*) test. The line and shading indicate the median and interquartile range, respectively. (**e**) Conservation of DNA sequence (measured by the PhastCons score of 100 vertebrate genomes) of eGenes and non-eGenes regarding tissue-sharing. The dot and bar represent mean and standard error, respectively. (**f**) Counts of four types of SNP-gene pairs across 34 tissues. Ind.: independent *cis*-eQTL. Top.: top *cis*-eQTL. Multi.: eGenes that have identical or high linkage disequilibrium (LD, *r^2^* > 0.8) *cis*-eQTL in any two tissues. Opp-multi.: eGenes that have an opposite direction of *cis*-eQTL effect between any two tissues. (**g**) Scatter plots of *cis*-eQTL effect sizes of 48 common multi-eGenes in blood and testis. *cis*-eQTL with the same directional effect are colored as blue (n = 36), and those with the opposite direction are colored as red (n = 12). (**h**) The *cis*-eQTL effects of *ODF2L* on chromosome 4 in blood and testis. Diamond symbols represent the top *cis*-eQTL of *ODF2L*.

We further found that an eGene tended to be regulated by *cis*-eQTL of smaller effect if it showed a higher level of tissue sharing or was expressed in more tissues (Fig. 3d, Extended Data Fig. 9f). The higher the tissue-sharing of eGenes, the larger the minor allele frequency (MAF) of their *cis*-eQTL, and the closer the distance of their *cis*-eQTL to TSS (Fig. 3d). This is in agreement with previous findings in GWAS applied to complex traits, i.e., genes with a higher level of pleiotropy tend to have smaller effects, presumable due in part to stronger underlying negative selection^7,22^. In addition, eGenes that were active in more tissues had a decreased PhastCons score (i.e., less sequence constraint), while genes that were not eGenes (non-eGenes) in more tissues had an increased PhastCons score (Fig. 3e). The shared non-eGenes (n = 1,511) in the 34 tissues were significantly enriched in fundamental biological processes, such as reproduction and organ development (Table S14). These findings suggest that genes without detectable *cis*-eQTLs in more tissues are more likely to be under stronger purifying selection due to their essential functions, resulting in a higher DNA sequence conservation across species.

We further examined whether eGenes exhibited the opposite direction of *cis*-eQTL effects between tissues. We summarized four types of SNP-gene pairs (Fig. 3f) and observed that 1.8% (1,166/64,250) of top *cis*-eQTL of the same eGenes had an opposite effect in at least one tissue pair, representing 3.1% (467/14,988) of all detected eGenes. Compared to other tissue pairs, blood and testis showed the highest proportion (25%) of eGenes with opposite *cis*-eQTL effects (Fig. 3g). For example, results for *ODF2L*, which participates in the negative regulation of cilium assembly, are shown in Fig. 3h. The top *cis*-eQTL (*rs329043485*) of *ODF2L* in blood was also the top *cis*-eQTL in testis, where the C allele of *rs329043485* increases *ODF2L* expression in blood but decreases expression of the same gene in testis (Fig. 3h, Extended Data Fig. 9g).

### Functional annotation of molQTL

All five types of molQTL were significantly enriched for missense and synonymous variants, variants in 3’ and 5’UTR, and variants with low and moderate effects on protein sequence. Compared to other molQTL, *cis*-sQTL had a higher enrichment for missense variants, variants with high impact on protein sequence, and variants in splice region and acceptor sites (Fig. 4a). Looking at chromatin states, these five types of molQTL showed the highest enrichment in active promoters, followed by those proximal to TSS and ATAC islands (Fig. 4b). In total, we found an average of 64% *cis*-eQTL could potentially modify transcription factor binding sites (TFBS) (Table S15). Although, on average, enhancers showed a weak enrichment of molQTL (Fig. 4b), we noted that enhancers had a higher enrichment for *cis*-eQTL in the matching tissue compared to non-matching tissues, demonstrating the important roles of enhancers and *cis*-eQTL on the tissue-specific regulation (Fig. 4c). Notably, the top *cis*-eQTL tended to be enriched in promoters rather than enhancers, whereas the reverse was observed for the second- and third-ranked *cis*-eQTL (Fig. 4d). This demonstrates that genetic variants in promoters generally had a larger effect on gene expression than those mediating enhancers. In addition, molQTL showed tissue-specific enrichment for hypomethylated regions (HMR) and allele specific methylation loci (ASM) (Fig. 4e). In muscle, through conducting MR analysis^23^, we observed that 2,016 *cis*-eQTL, 4,694 *cis*-eeQTL, 524 *cis*-lncQTL, 5,174 *cis*-enQTL and 1,590 *cis*-sQTL were mediated by meQTL (Fig. S11, Table S16). For instance, we found that the methylation level of a CpG locus on chromosome 16 was significantly associated with the expression of *DHX29*, and also associated with a complex trait, i.e., loin muscle area (LMA) (Fig. 4f). By overlapping *cis*-eQTL with topologically associating domain (TAD), we found that long-distance *cis*-eQTL were significantly enriched for the same TAD as TSS of target genes after accounting for the *cis*-eQTL-TSS distance, which was consistent across tissues (Fig. 4g). This suggests that long-range *cis*-eQTL may affect gene expression by mediating physical interactions of the chromosome regions^24^. We took *BUD23* in muscle as an example (Fig. 4h). The second independent *cis*-eQTL of *BUD23* was 385kb upstream of its TSS, and located within the same TAD of the TSS, as well as was surrounded by HMRs and enhancers in muscle. Altogether, the results here indicate the complexity of the molecular mechanisms underlying these regulatory variants on target gene expression.

**Figure 4.**
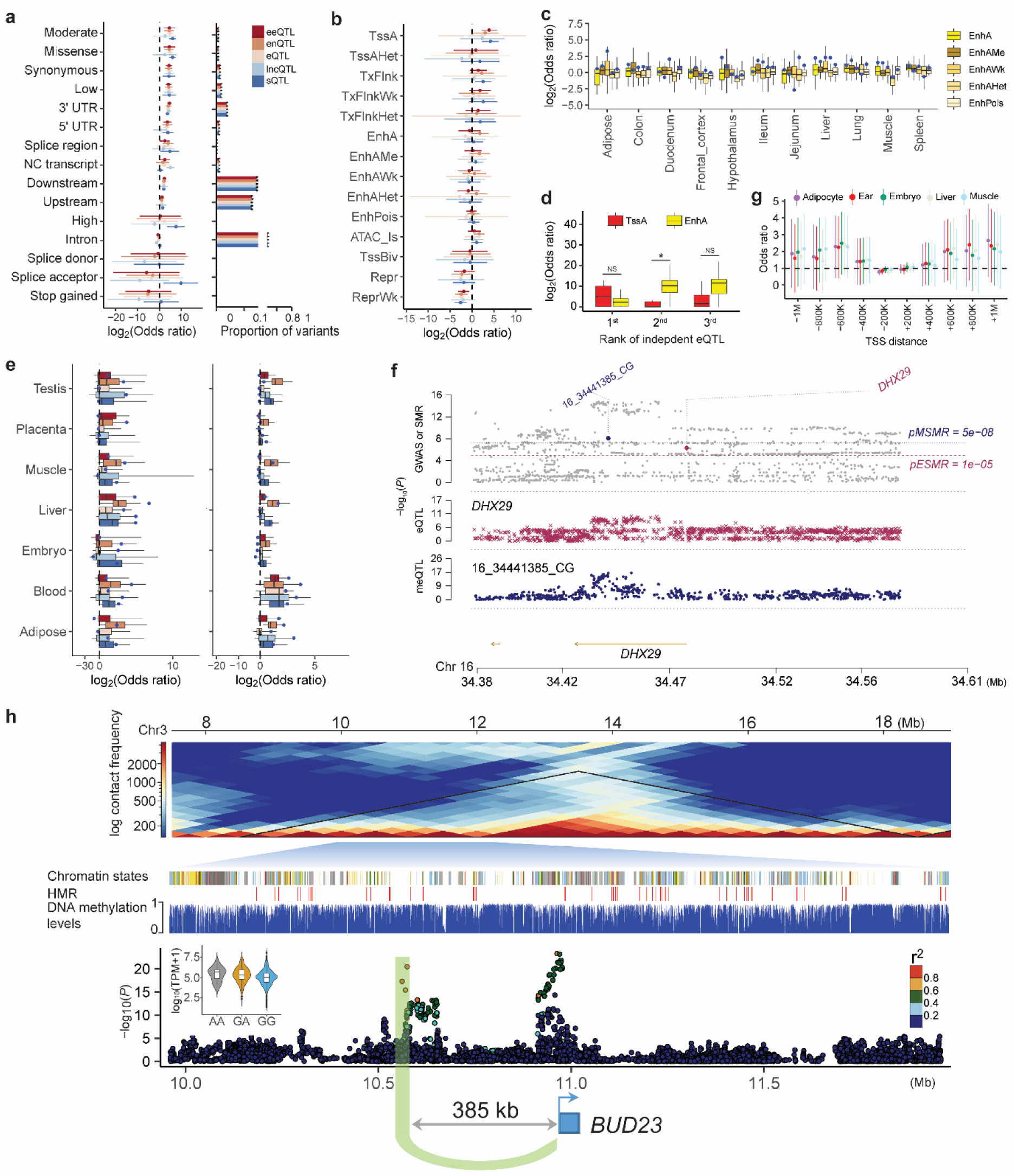
Functional characterization of regulatory variants. Enrichment (odds ratio) of five types of molQTL in sequence ontology (**a**) and 14 chromatin states^17^ (**b**). Enrichment is shown as mean ± standard deviation across 34 tissues. (**c**) Enrichment of *cis*-eQTL in five types of enhancers. Each box includes enrichment of *cis*-eQTL from 34 tissues across enhancers. Blue dots represent enrichments from matching tissues. (**d**) Enrichment of top three independent *cis*-eQTL in two chromatin states. TssA is for active transcription start sites (TSS), while EnhA is for active enhancers. The *P* values are obtained by the two-sided Student *t*-test. * indicates *P* < 0.05 and NS indicates not significant. (**e**) Enrichment of five molQTL types in DNA hypomethylated regions (HMR, left) and allele-specific methylation loci (ASM, right) across seven tissues that have both DNA methylation and *cis*-eQTL data. The color key of molQTL is the same as in (**a**). Blue dots represent enrichments from methylation-molQTL matching tissues. (**f**) The summary data-based Mendelian randomization (SMR) results of methylation QTL (meQTL), *cis*-eQTL and GWAS of loin muscle area around *DHX29* gene in muscle. The top plot shows −log_10_(*P*) of SNPs from GWAS. The red diamond and blue circle represent −log_10_(*P*) from the SMR test for associations with gene expression and DNA methylation, respectively. The middle plot shows *cis*-eQTL results of *DHX29*. The bottom plot shows meQTL results at the CpG locus (16_34441285_CG). (**g**) Enrichment of *cis*-eQTL within the same topologically associating domain (TAD) of TSS of target genes. TADs are obtained from Hi-C data of five tissues. The *cis*-eQTL are grouped according to their distance to TSS. – and + means upstream and downstream, respectively. (**h**) The landscape of *BUD23* at multiple genomic features in muscle. The top plot shows that *BUD23* and its second independent eVariant (rs790620973) are located within a TAD. The bottom is the Manhattan plot showing *cis*-eQTL results of *BUD23*. The violin plot shows expression levels of *BUD23* across three genotypes of this eVariant in muscle. There are 9, 131, and 1,181 samples for AA, GA, and GG genotypes, respectively. TPM: Transcripts Per Million.

### Interpreting GWAS loci with molQTL

To study the regulatory mechanisms underlying complex traits in pigs, we examined 268 meta-GWAS, representing 207 distinct traits belonging to five categories, including reproduction traits (n = 71), health traits (n = 61), meat and carcass traits (n = 50), production traits (n = 19) and exterior traits (n = 6) (Table S17). Enrichment analysis showed that GWAS signals of all traits were significantly enriched in all five types of molQTL (Fig. 5a, Fig. S12a-e). Among them, *cis*-eQTL and *cis*-sQTL showed the highest enrichment (~1.61-fold, *se* =0.014), followed by *cis*-eeQTL (1.57-fold, *se* = 0.015), *cis*-enQTL (1.56-fold, *se* = 0.014), and *cis*-lncQTL (1.55-fold, *se* = 0.014) (Fig. 5a, Fig. S12f). Subsequently, we quantified the proportion of heritability mediated by these molecular phenotypes using MESC^25^ for 198 complex traits with enough data. Averaging across traits, we found that approximately half of the heritability was mediated by PCG expression and alternative splicing, followed by enhancer expression (48.3%), exon expression (46.4%), and lncRNA expression (29.1%) (Fig. 5b). These results are comparable to our previous findings in cattle where e/sQTL from 16 tissues explained approximately 70% of the heritability of complex traits^26^.

**Figure 5.**
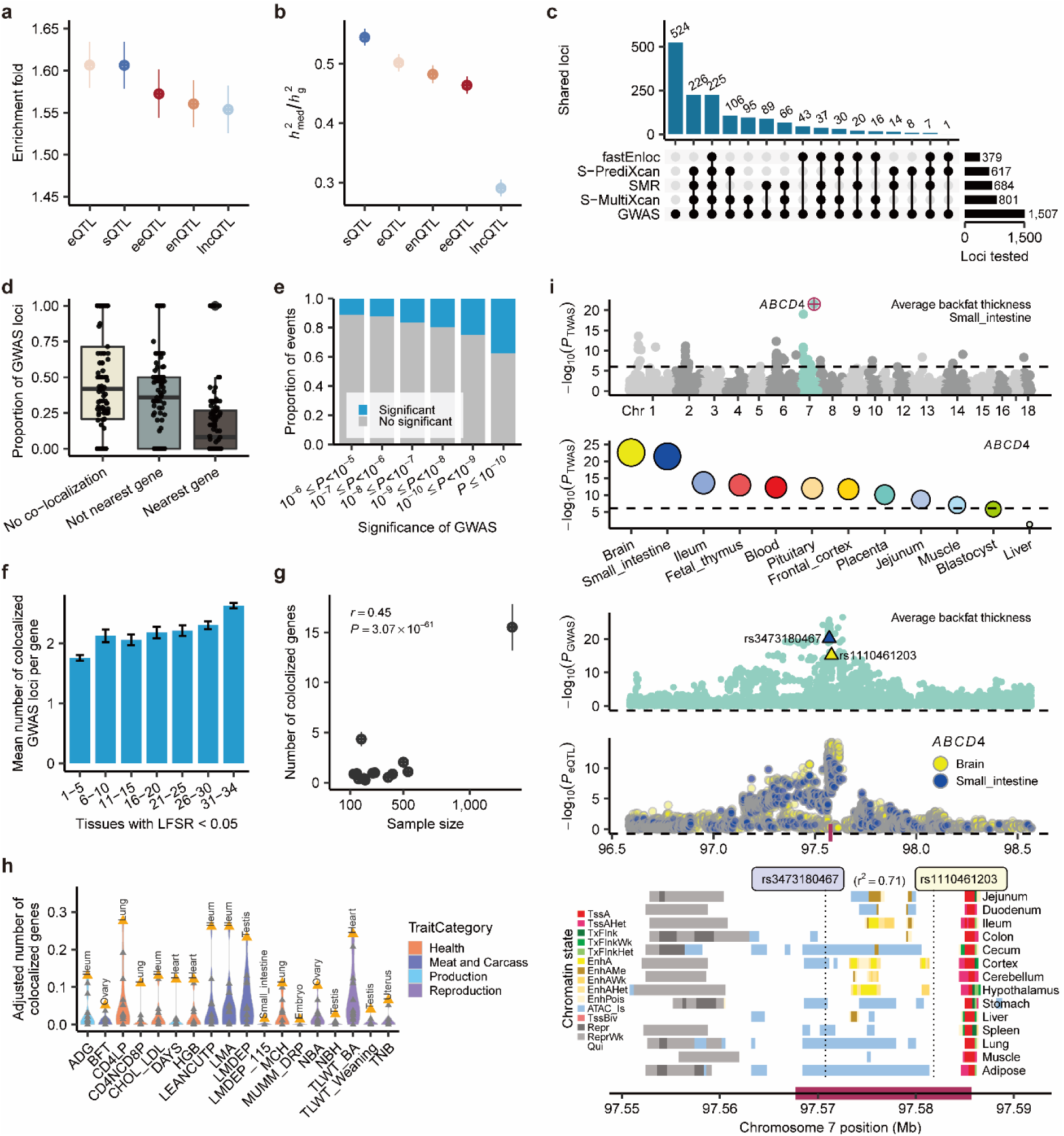
Interpreting GWAS loci of complex traits using molecular QTL (molQTL). (**a**) Enrichment (mean and 95% confidence interval, CI) of GWAS variants (*P* ≤ 0.05) with five types of molQTL in 34 tissues. We consider 198 GWAS with ≥ 80% of variants being tested in the molQTL mapping. (**b**) Proportion (mean and 95% CI) of heritability (ℎ^2^) mediated by the *cis*-genetic component of five molecular phenotypes (ℎ^2^) in 34 tissues across GWAS traits. (**c**) Number of GWAS loci linked to eGenes in 34 tissues through fastEnloc (colocalization), SMR (Mendelian randomization), S-PrediXcan (single-tissue transcriptome-wide association studies, TWAS) and S-MultiXcan (multi-tissue TWAS). (**d**) Proportion of three types of GWAS loci regarding the colocalization results. No colocalization: GWAS loci that are not colocalized with any eGenes in 34 tissues. Not nearest gene: GWAS loci whose colocalized eGenes are not nearest genes to GWAS lead SNPs. Nearest gene: GWAS loci whose colocalized eGenes are nearest ones. Each dot represents a complex trait. We consider 105 GWAS traits with significant colocalizations (i.e., regional colocalization probability, RCP > 0.9). (**e**) Proportion of significant colocalizations of *cis*-eQTL with GWAS loci at different significance levels. (**f**) The mean number of colocalized GWAS loci per eGene across 105 traits above. eGenes are classified into seven groups regarding the tissue-sharing pattern. Error bar indicates the standard error. (**g**) Relationship between the number of colocalized genes in GWAS traits and *cis*-eQTL tissue sample size. Point and error bar indicate the mean and standard error, respectively. We only consider 14 tissues with over 100 samples and 91 GWAS traits with RCP > 0.9. (**h**) The number of colocalized genes adjusted for tissue sample size and eGene discovery ratio in 14 tissues across 18 GWAS traits. Top tissues are labeled. The detailed definitions of traits are summarized in Table S17. (**i**) The association of *ABCD4* with average backfat thickness (BFT). The top Manhattan plot represents the single-tissue TWAS results of BFT in small intestine. Followed by the single-tissue TWAS results of *ABCD4* for BFT in 12 tissues being tested. The two following Manhattan plots show the colocalization of BFT GWAS (up) and *cis*-eQTL (down) of *ABCD4* on chromosome 7 in both brain and small intestine. The blue and yellow triangles indicate the colocalized variants of *ABCD4* in small intestine (rs3473180467) and brain (rs1110461203), respectively. These two variants have a high linkage disequilibrium (LD) (*r*^2^ = 0.71). The bottom panel is for chromatin states around *ABCD4* on chromosome 7.

Although the above enrichment analysis demonstrated the overall enrichment of GWAS signals with molQTL, it was limited to interpreting the regulatory mechanisms underlying individual significant GWAS loci. Therefore, we employed four complementary approaches to detect shared regulatory variants/genes associated with both molecular phenotypes and complex traits, including colocalization *via* fastENLOC^27^, Mendelian randomization *via* SMR^23^, single-tissue TWAS *via* S-PrediXcan^28^, and multi-tissue TWAS *via* S-MultiXcan^29^, and all referred to as ‘colocalization’ hereafter. Here we only considered 1,507 significant GWAS loci that were tested in the *cis*-eQTL mapping. Out of these, 983 (65%) GWAS loci were colocalized with *cis*-eQTL in at least one tissue tested (Fig. 5c, Table S18). Among these 983 colocalization events, only 33% were colocalized with the nearest genes of the lead GWAS SNP (Fig. 5d), which indicates the regulatory complexity of complex traits. We observed that GWAS loci mapped with higher confidence level were more likely to be colocalized with *cis*-eQTL (Fig. 5e). The eGenes shared between more tissues tended to be colocalized with more GWAS loci across complex traits (Fig. 5f). Of note, the number of colocalization events of a trait was determined by the statistical power of both GWAS and *cis*-eQTL mapping (Fig. 5g, Fig. S12g-h, Fig. S13).

To prioritize tissues relevant for complex trait variation, we defined a ‘tissue relevance score’ through the number of colocalization events adjusted by sample size and eGene discovery ratio of a tissue (Table S19). We only included 14 tissues with over 100 samples due to needed statistical power for *cis*-eQTL mapping. We found that, for instance, ileum was the most relevant tissue for both average daily gain (ADG) and loin muscle area (LMA), and uterus for total number of piglets born (TNB) (Fig. 5h). We highlight *ABCD4* as an example (Fig. 5i) to illustrate how the PigGTEx resource can improve our understanding of the genetic regulatory mechanism underpinning complex traits in pigs. First, *ABCD4* was the top associated gene (*P*_TWAS_=3.18×10^−22^) in the small intestine TWAS of average backfat thickness (BFT). *ABCD4* also had a significant association with BFT in other tissues, where brain was the most significant, followed by ileum, fetal thymus, and blood. We then observed a significant colocalization between GWAS loci of BFT and *cis*-eQTL of *ABCD4* in both brain and small intestine. Although the top colocalized SNPs were different in brain (rs1110461203) and in small intestine (rs3473180467), they had a relatively high LD (*r*^2^ = 0.71), potentially tagging the same underlying causal variant. We found active enhancers around these two colocalized SNPs in brain and intestine but not in other tissues. ABCD4 participates in fatty acid catabolic processes and the transportation of vitamin B_12_ across liposomal membranes^30^.

Furthermore, we employed the same GWAS integrative analysis for other molQTL (Table S20-22). In total, we found that ~80% (1,197/1,507) of significant GWAS loci could be explained by at least one molQTL across the 34 tissues. Of note, 8.2%, 3.8%, 3.5% and 1.9% of all 1,507 GWAS loci were only explained by *cis*-eQTL, *cis*-sQTL, *cis*-eeQTL and *cis-*lncQTL, respectively, indicating that each molecular phenotype can make a unique contribution to biologically interpreting complex traits at distinct levels of gene regulation (Extended Data Fig. 10a-b). For example, a GWAS signal of ADG on chromosome 13 was only significantly colocalized with *cis*-eQTL of *CFAP298-TCP10L* in colon, but not with its *cis*-sQTL or *cis*-eeQTL (Extended Data Fig. 10c). The GWAS signal for BFT on chromosome 15 was exclusively colocalized with *cis*-sQTL of *MYO7B* in small intestine, while the GWAS signal of litter weight was exclusively colocalized with *cis*-eeQTL of *FBXL12* in uterus (Extended Data Fig. 10d-e). In addition, the multi-layer data of PigGTEx provides the opportunity to understand the etiology of genetic control of complex traits. We thus performed MR analysis to integrate GWAS with both *cis*-lncQTL and *cis*-eQTL, and detected 512 lncRNA-PCG-trait trios with significant pleotropic associations (Table S23). For instance, a *cis*-eQTL of a lncRNA (*MSTRG.4694&ENSSSCT00000070888*), residing downstream of *GOSR2*, had a significant colocalization with a *cis*-eQTL of *GOSR2* in muscle and GWAS signals of loin muscle depth (LMD) on chromosome 12 (Extended Data Fig. 10f). This implies a plausible biological pathway indicating that lncRNA *MSTRG.4694&ENSSSCT00000070888* regulates *GOSR2* expression in muscle, leading to changes in LMD in pigs. All these findings here demonstrated that PigGTEx provides a highly valuable resource for interpreting the biological and regulatory basis of complex traits in pigs.

### Do pigs share gene regulation and genetics of complex phenotype with humans?

By examining GTEx (v8) in humans^7^, we first compared gene expression across 17 common tissues between humans and pigs, including 8,540 and 3,913 samples in humans and pigs, respectively. We found that one-to-one orthologous genes (n = 15,944, 95.56% were PCGs) contributed to an average of 82% and 87% of transcriptional outputs across tissues in pigs and humans, respectively. This was consistent between species (Pearson’s *r* = 0.75, *P* = 5.21 × 10^−4^) (Fig. S14a-b). Compared to other tissues, orthologous genes in blood contributed the smallest proportion of transcriptional outputs in both species. The visualization of variation in gene expression among all 12,453 samples clearly recapitulated tissue types rather than species (Fig. S14c), demonstrating the overall conservation of gene expression between humans and pigs (Fig. 6a, Fig. S14d). We further found that genes with high or low expression were more conserved than those with moderate expression (Fig. S14e), and tissue-specific expressed genes were more conserved than the rest (Fig. S14f-g).

**Figure 6.**
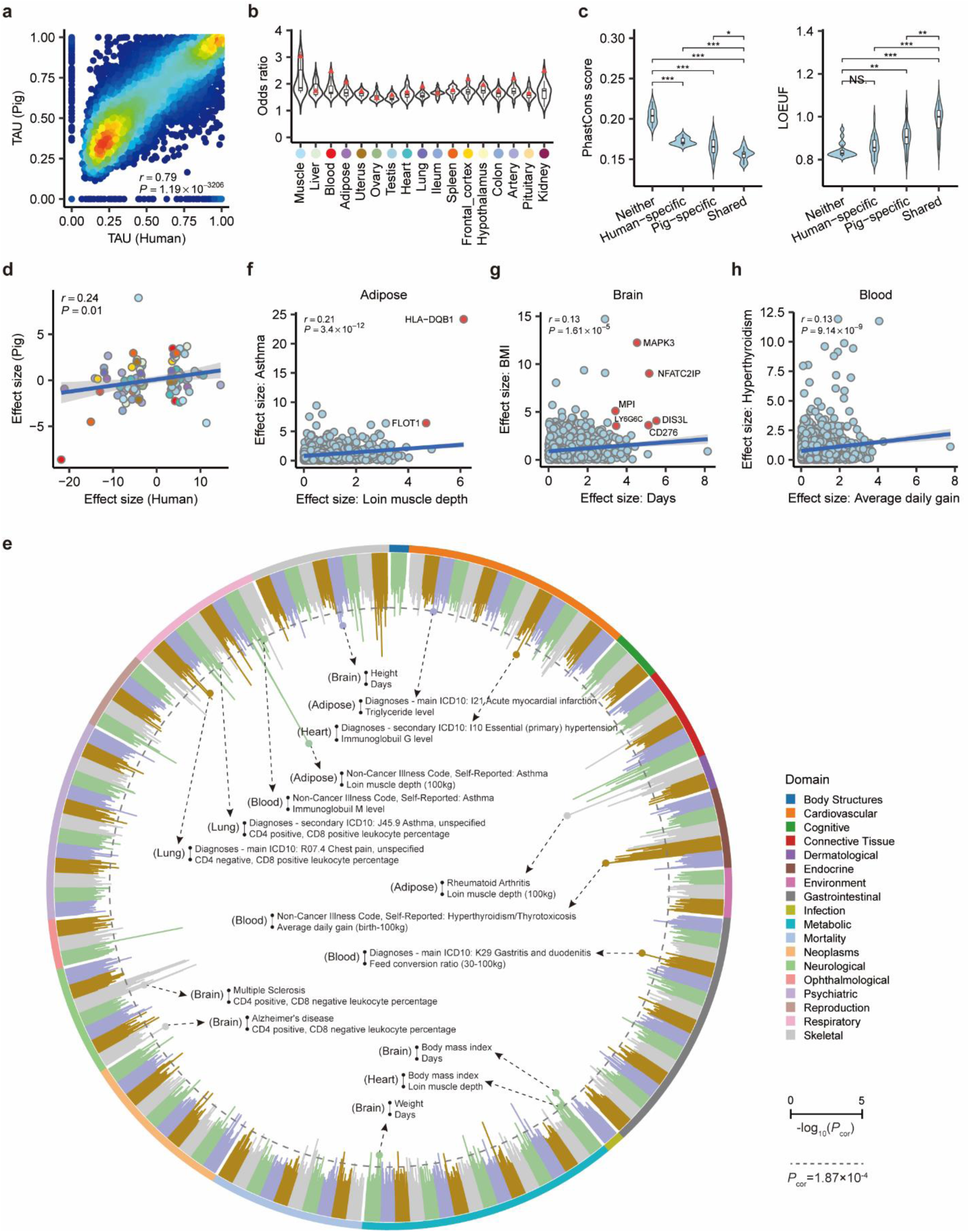
Conservation of gene expression, *cis*-eQTL and complex trait genetics between pigs and humans. (**a**) Pearson’s correlation (*r*) of TAU values (measuring the tissue-specificity of gene expression) of 14,843 one-to-one orthologous genes between humans and pigs. Dots represent genes, and colors show the density of dots. (**b**) Enrichment (odds ratio, obtained by Fisher’s exact test) of pig eGenes with human eGenes across 17 shared tissues. Red points represent enrichments in matching tissues. (**c**) Comparison of DNA sequence conservation and tolerance to loss of function mutations (LOEUF) between four groups of orthologous genes across 17 tissues, including non-eGenes in neither species (Neither, n = 15,750), human-specific eGenes (Human-specific, n = 14,394), pig-specific eGenes (Pig-specific, n = 9,038) and eGenes shared in both species (Shared, n = 9,731). The two-sided Wilcoxon test is used to test the difference between groups. NS., *, **, and *** represent non-significant, *P* < 0.05, 0.01, and 0.001, respectively. (**d**) Pearson’s *r* of effects of 112 orthologous variants on gene expression between humans and pigs. Each point represents a variant, which is a significant *cis*-eQTL in humans. Dot color represents tissue type, same as in (**b**). (**e**) Significance (-log_10_*P*) of Pearson’s *r* of orthologous gene effect size between pig and human traits derived from transcriptome-wide association studies (TWAS). Each bar represents the −log_10_(*P*) of a pig-human trait pair in the same tissue. Labels in the middle are significant examples of pig-human trait pairs (*P* < 1.87×10^−4^, based on the Bonferroni correction), including the human trait (top), the pig trait (bottom), and the TWAS tissue (left). (**f**) Scatter plot for effect sizes of 1,121 orthologous genes between loin muscle depth (pig) and asthma (human) traits in adipose. (**g**) Effect sizes of 1,177 orthologous genes between days (pig) and BMI (human) in brain. (**h**) Effect sizes of 1,872 orthologous genes between average daily gain (pig) and hyperthyroidism (human) in blood. Dots represent orthologous genes. Red dots represent genes with significant (FDR < 0.05) TWAS signals in both species. The lines are fitted by a linear regression model.

By comparing tissue-sharing patterns of eGenes between pigs and humans, we found that the number of tissues in which an eGene was active was significantly correlated between species (Fig. S14h). We found a total of 136 eGenes that were active in more than 10 tissues in both species, which were significantly enriched in metabolic processes like organic acid catabolic process (Table S24). A total of 999 genes were not eGenes in any tissues neither species, and these were significantly enriched in the essential biological functions such as sensory perception of chemical stimulus and nervous system process (Table S25). We further observed that eGenes in a pig tissue generally had a higher enrichment for eGenes in the matching tissue in humans compared to other tissues (Fig. 6b), even though the tissue sample sizes were different between species. Compared to other genes, eGenes shared by both species had the lowest evolutionary DNA sequence conservation and the highest tolerance to inactivation, whereas shared non-eGenes showed the opposite trend, indicative of their essential functions and underlying strong purifying selection (Fig. 6c). Furthermore, we found that 112 *cis*-eQTLs in humans had orthologous variants in pigs, and their effects on gene expression were significantly correlated between the species (Fig. 6d, Table S26). All these findings together provided evidence that the genetic regulation of gene expression is conserved to a certain degree between pigs and humans.

To investigate whether the regulatory mechanism of complex phenotype was conserved between humans and pigs, we systematically compared effect sizes of orthologous genes between 268 pig and 136 human complex traits and diseases based on the summary statistics of TWAS (Table S27). We found a total of 340 significant (Bonferroni corrected-*P* < 0.05) pig-human trait pairs within the same tissue (Table S28, Fig. 6e). For instance, the most significant correlation (Pearson’s *r* =0.21, *P*-value=3.4×10^−12^) was observed for human asthma and pig LMD in adipose, and *HLA-DQB1* was the top associated gene in both species (Fig. 6f). This is consistent with previous findings that asthma is associated with obesity/adipose and smooth muscle cells in humans^31,32^. We also found significant correlations for human BMI *vs*. pig growth (i.e., days to 100kg) in brain and human hyperthyroidism *vs*. pig ADG in blood (Fig. 6g-h). These findings are supported by previous studies suggesting that rapid growth is associated with BMI in humans^33^, and hyperthyroidism has effects on growth and development in pigs^34,35^. All these findings corroborate the use of the pig as a biomedical model for certain human diseases and complex traits, particularly those related with fat deposition and growth.

## Discussion

As an essential part of the FarmGTEx project, the pilot phase of PigGTEx offers a deep survey of gene expression and regulatory effects across a wide range of tissues, representing a massive escalation in understanding of the gene regulation landscape in pigs. Compared to the previous gene expression atlas based on a smaller number of individuals^36,37^, this large gene expression atlas enables us to better annotate the function of genes in pigs through investigating tissue-specific expression and co-expression patterns. The multi-tissue catalogue of regulatory variants is a prerequisite for functional annotation and thus, advances our understanding of biological mechanisms underlying complex traits of economic importance and animal welfare in pigs. On average, about 80% of GWAS loci tested are explained by candidate genes in pigs, comparable to 78% of GWAS loci deciphered by GTEx in humans^7^. The established resource will eventually enhance genetic improvement programs through the development of advanced biology-driven genomic prediction models that depend on informative SNPs ^38^. We also demonstrate the level of similarity between pigs and humans in gene expression, regulatory effects and complex traits genetics. This extensive comparison of the pig and human genomes at multiple biological levels will be instructive for deciding on which human diseases and complex traits the pig is the most suitable animal model for.

Characterizing the tissue specificity of regulatory effects is of utmost importance in identifying the causal tissues and dissecting molecular mechanisms underlying complex traits of economic importance in pigs. Although a fraction of regulatory effects is shared across tissues, we note that some tissues like testis and those from early developmental stages are distinct from other primary tissues. To gain a more comprehensive view of tissue-specific gene regulation and refine tissue-trait map, underrepresented tissues (like different brain regions, mammary, osseous and epithelial tissues) at multiple development stages are still required in pigs. To elucidate gene regulation at single cell resolution, we conducted an exploratory analysis to discover cell type-interaction regulatory effects through an *in silico* cell type deconvolution^21^. The cieQTL identified for several cell types indicate that a vast majority of cell type–specific *cis*-QTL remains to be detected^39,40^. A multi-tissue pig cell atlas has recently been made available^41^, which will provide greater precision in cell type deconvolution in future studies. Here, we do not consider *trans*-eQTL (> 1Mb to the TSS of genes) due to the current small sample size across tissues. Indeed, compared to cis-eQTL, trans-eQTL often have smaller effect sizes and tend to be non-additive inherited among many species, and thus require hundreds of thousands of samples and more comprehensive GWAS models to be discovered^24,42^. Although integrating multi-omics data provides insight into the molecular mechanisms underlying regulatory variants, experimental follow-ups (massively parallel reporter assays and genome-wide CRISPR perturbations) are necessary to functionally validate and characterize these regulatory variants at large scale^43,44^.

## Acknowledgements

Z. Zhang (SCAU) acknowledges funding from the National Natural Science Foundation of China (32022078), the Local Innovative and Research Teams Project of Guangdong Province (2019BT02N630), and supporting from National Supercomputer Center in Guangzhou China. Y. Chen, Z. Zhang (SCAU), Jiaqi Li, X. Liu, X. Ding, and S. Zhao acknowledge funding from the China Agriculture Research System (CARS-35). L. Fang. acknowledges funding from HDR-UK award HDR-9004 and the European Union’s Horizon 2020 research and innovation program under the Marie Skłodowska-Curie grant agreement No 801215. G.E.L., was supported by USDA NIFA AFRI grant numbers 2019-67015-29321 and 2021-67015-33409 and the appropriated project 8042-31000-112-00-D, “Accelerating Genetic Improvement of Ruminants Through Enhanced Genome Assembly, Annotation, and Selection” of the USDA Agricultural Research Service (ARS). This research used resources provided by the SCINet project of the USDA ARS project number 0500-00093-001-00-D. Mention of trade names or commercial products in this article is solely for the purpose of providing specific information and does not imply recommendation or endorsement by the USDA. The USDA is an equal opportunity provider and employer. A. Tenesa acknowledges funding from the BBSRC through programme grants BBS/E/D/10002070 and BBS/E/D/30002275, MRC research grant MR/P015514/1, and HDR-UK award HDR-9004. P. Navarro and C. Haley were supported by the Medical Research Council (United Kingdom, grant number MC_UU_00007/10). O. Canela-Xandri was supported by MR/R025851/1. M. Ballester and D. Crespo-Piazuelo belonged to a Consolidated Research Group AGAUR, ref. 2017SGR-1719, and D. Crespo-Piazuelo was supported by the GENE-SWitCH project (https://www.gene-switch.eu), which is funded by the European Union’s Horizon 2020 research and innovation programme under the grant agreement n° 817998. R.X. was supported by Australian Research Council’s Discovery Projects (DP200100499). L.M. was supported in part by AFRI grant numbers 2020-67015-31398 and 2021-67015-33409 from the United States Department of Agriculture (USDA) National Institute of Food and Agriculture (NIFA). B.N.K and G.A.R. were supported by appropriated project 3040-31000-099-000-D, “Identifying Genomic Solutions to Improve Efficiency of Swine Production” of the Agricultural Research Service (ARS) of the United States Department of Agriculture (USDA). A.K.L.P and W.T.O were supported by appropriated project 3040-31000-102-000-D, “Optimizing Nutrient Management and Efficiency of Beef Cattle and Swine” of the Agricultural Research Service (ARS) of the United States Department of Agriculture (USDA). Z.P, D.G, and H.Zhou, and computational resource was supported in part by Agriculture and Food Research Initiative Competitive Grants no. 2018-67015-27501 and no. 2015-67015-22940. All the funders had no role in study design, data collection and analysis, decision to publish or preparation of the manuscript.

We thank all the researchers who have contributed to the publicly available data used in this research. We thank the valuable comments and suggestions from Prof. Doug Speed, Dr. Guillaume Paul Ramstein (QGG, Aarhus University, Denmark), Prof. Michael E Goddard (The University of Melbourne, Australia), Prof. Chris Ponting (IGC, The University of Edinburgh, UK), and Prof. Greger Larson (The University of Oxford, UK). Figure 1d was created with BioRender.com. For the purpose of open access, the author has applied a Creative Commons Attribution (CC BY) license to any Author Accepted Manuscript version arising from this submission.

## Author Contributions Statement

L.Fang, Z.Zhang (SCAU), G.E.L., A.T., and K.L. conceived and designed the project. Y.G., S.L., X.Li, H.Y., B.Z., W.Yang, W.Yao, Y.Y., H.L., H.Zhang., and X.P. performed bioinformatic analyses of RNA-Seq data analysis. H.Y., S.D., L.B., S.W., D.G., L.Y., and Z.Chen. conducted whole-genome sequence data analysis. Y.G., Q.Zhao, and Z.P., performed omics data analysis. J.T., conducted genotype imputation and molQTL mapping. Z.X., H.Zeng, C.W., W.L., T.C., and X.Yu, prepared the summary statistics of GWAS in pigs and humans. J.T., Q.L., X.C., and J.W., integrated molQTL with GWAS. Z.B., J.T., C.X., and Jinghui Li led the comparison of PigGTEx and human GTEx. B.N.K., G.A.R., A.K.L.P., W.T.O., M.B., D.C., M.C., and L.K., contributed to the validation and functional annotation of molQTL. P.N., Y.H., B.L., Z.Cai, P.Z., D.R., C.L., H.P., X.H., L.Frantz, Y.L., L.L., L.C., J.J., R.H., Z.T,. M.L., S.Z., and Y.C., contributed to the critical interpretation of analytical results before and during manuscript preparation. H.Zeng, J.T., Z.Zhang (SCAU) and L.Fang., built the PigGTEx web portal. L.Fang, Z.Zhang (SCAU), G.E.L., K.L., M.B., R.Q., O.C.-X., K.R., P.K.M, M.F., M.A., A.C., E.G., H.C., G.Su, G.Sahana, M.S.L., J.C.M.D., C.K.T., R.C., M.A.M.G., O.M., M.G., Z.Zhou, Z.Zhang (ZJU), R.X., X.S., P.L., G.T., Y.Z., G.Y., F.Z., P.N., X.Yuan, X.Liu, L.M., H.S., X.X., Q.W., X.D., H.Zhou, Jiaqi Li, C.H., Y.P., B.L., and Q.Zhang, contributed to the data and computational resources. L.Fang, J.T., Y.G., and Z.B., drafted the manuscript. All authors read, edited, and approved the final manuscript.

## Extended Data Figures and legends

**Extended Data Figure 1.**
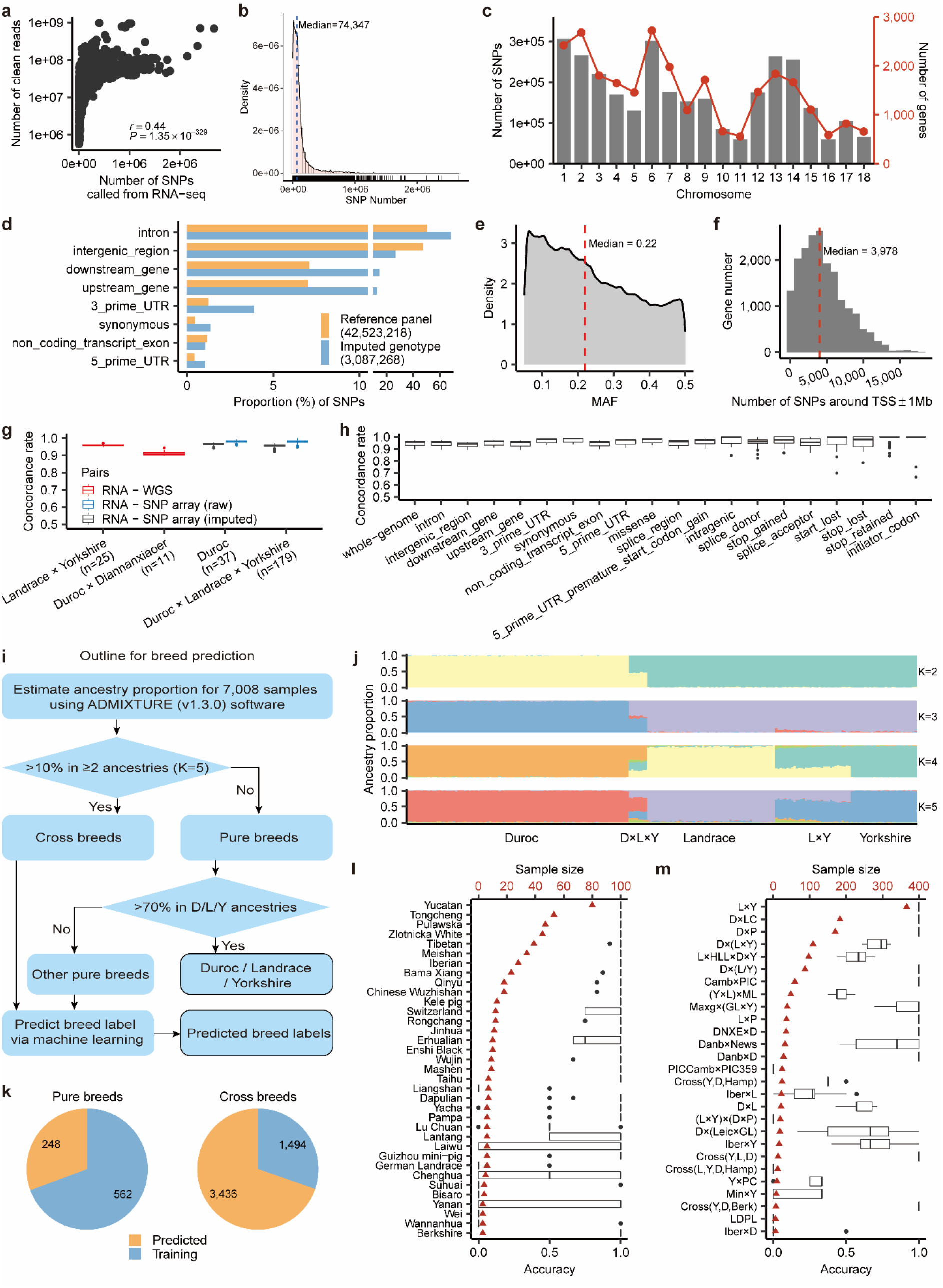
Genotype calling and imputation and breed prediction. (**a**) Pearson’s correlation (*r*) between number of clean reads and number of called SNPs across 7,095 RNA-Seq samples. The *P*-value is obtained by Pearson’s *r* test. (**b**) Distribution of number of SNPs called from 7,095 RNA-Seq samples. (**c**) Number of imputed SNPs (left, gray bars) from 7,008 RNA-Seq samples across 18 pig chromosomes after quality control (DR^2^ ≥ 0.85, minor allele frequency ≥ 0.05). The red point represents number of genes (right) in each chromosome in the Sscrofa11.1. assembly (Ensembl v100). (**d**) Distribution of 42,523,218 SNPs from the Pig Genomics Reference Panel (PGRP) and 3,087,268 imputed SNPs used for molecular QTL (molQTL) mapping across eight genomic regions annotated by SnpEff (v.4.3). (**e**) Minor allele frequency (MAF) of imputed SNPs across 7,008 RNA-Seq samples. (**f**) Distribution of number of imputed SNPs around 1Mb of transcript start site (TSS) of 18,911 protein-coding genes. (**g**) Concordance rates between imputed genotypes from RNA-Seq only and those directly called from whole-genome sequence (WGS) data (red), raw genotypes (blue) from SNP array and imputed genotypes (gray) from SNP array, respectively, in the same individuals. Labels of *x*-axis are breeds and number of individuals. (**h**) Imputation accuracy (concordance rate) of SNPs across different genomic regions annotated by SnpEff (v.4.3). (**i**) The overall pipeline utilized to predict missing breed labels for RNA-Seq samples. (**j**) Estimated ancestry proportion of Duroc (n = 485), Landrace (n = 280), Yorkshire (n = 145), Landrace×Yorkshire (n = 165) and Duroc×Landrace×Yorkshire (n = 40) samples using ADMIXTURE (v1.3.0). (**k**) Distribution of sample size of training and prediction sets in pure and cross breeds. (**l-m**) Accuracy of breed prediction for pure breeds (**l**) and cross breeds (**m**) measured by 10 times three-fold cross-validation. Red triangle represents the sample size of target breed.

**Extended Data Figure 2.**
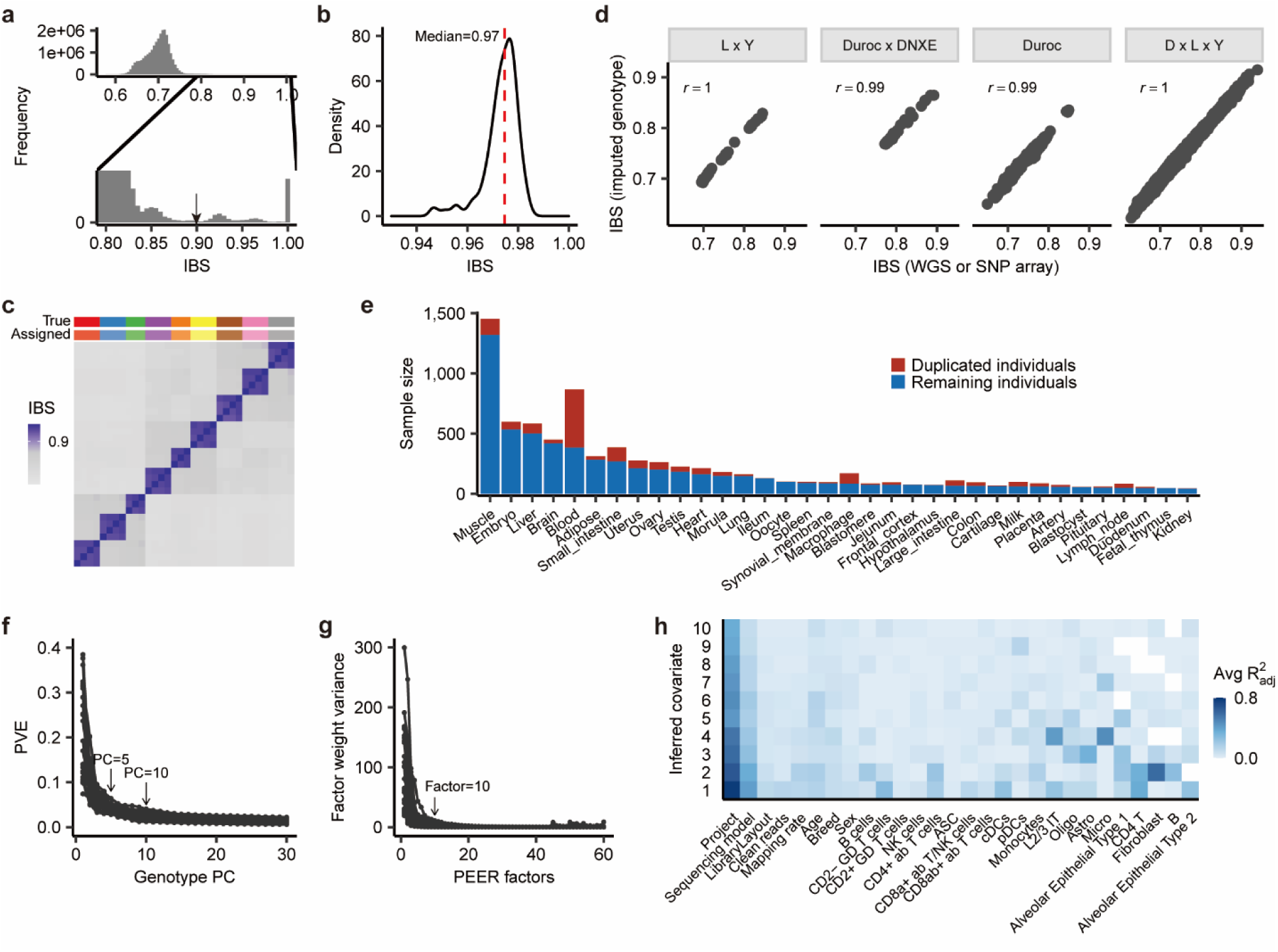
Detection of duplicated individuals and confounders of RNA-Seq samples. (**a**) Distribution of identity-by-state (IBS) distances among 7,008 RNA-Seq samples, which are calculated using 12,207 LD-independent SNPs (*r*^2^ < 0.2). (**b**) Density of IBS distances that are computed using genotypes derived from RNA-Seq only and whole-genome sequence (WGS) or SNP array data in the same individuals (n = 227). (**c**) Heatmap of IBS distance of 25 RNA-Seq samples from 9 individuals. The same color on the top of panel (**c**) represents samples from same individuals. True: true individual label; Assigned: assigned individual label using an IBS distance cutoff of 0.9. (**d**) Pearson’s correlation (*r*) between IBS distance calculated from imputed genotypes and those calculated from WGS or SNP array data across four different populations. L × Y: Landrace and Yorkshire cross breed (n = 25); Duroc × DNXE: Duroc and Diannanxiaoer cross breed (n = 11); Duroc: Duroc pure breed (n = 37); D × L × Y: composite population with 1/3 Duroc, 1/3 Landrace and 1/3 Yorkshire (n = 179). (**e**) Duplicated and remaining (unique) individuals in each of 34 pig tissues used for molecular QTL mapping. Sample pairs with IBS > 0.9 are considered as duplicated individuals. (**f**) Proportion of variance explained (PVE) by genotype principal components (PC) in each of 34 tissues (lines). (**g**) Factor weight variance of Probabilistic Estimation of Expression Residuals (PEER) factors in each of 34 tissues (lines). (**h**) Proportion of variance (adjusted R^2^) of known confounders captured by the top 10 inferred PEER factors, calculated using the *lm* function in R (v4.0.2).

**Extended Data Figure 3.**
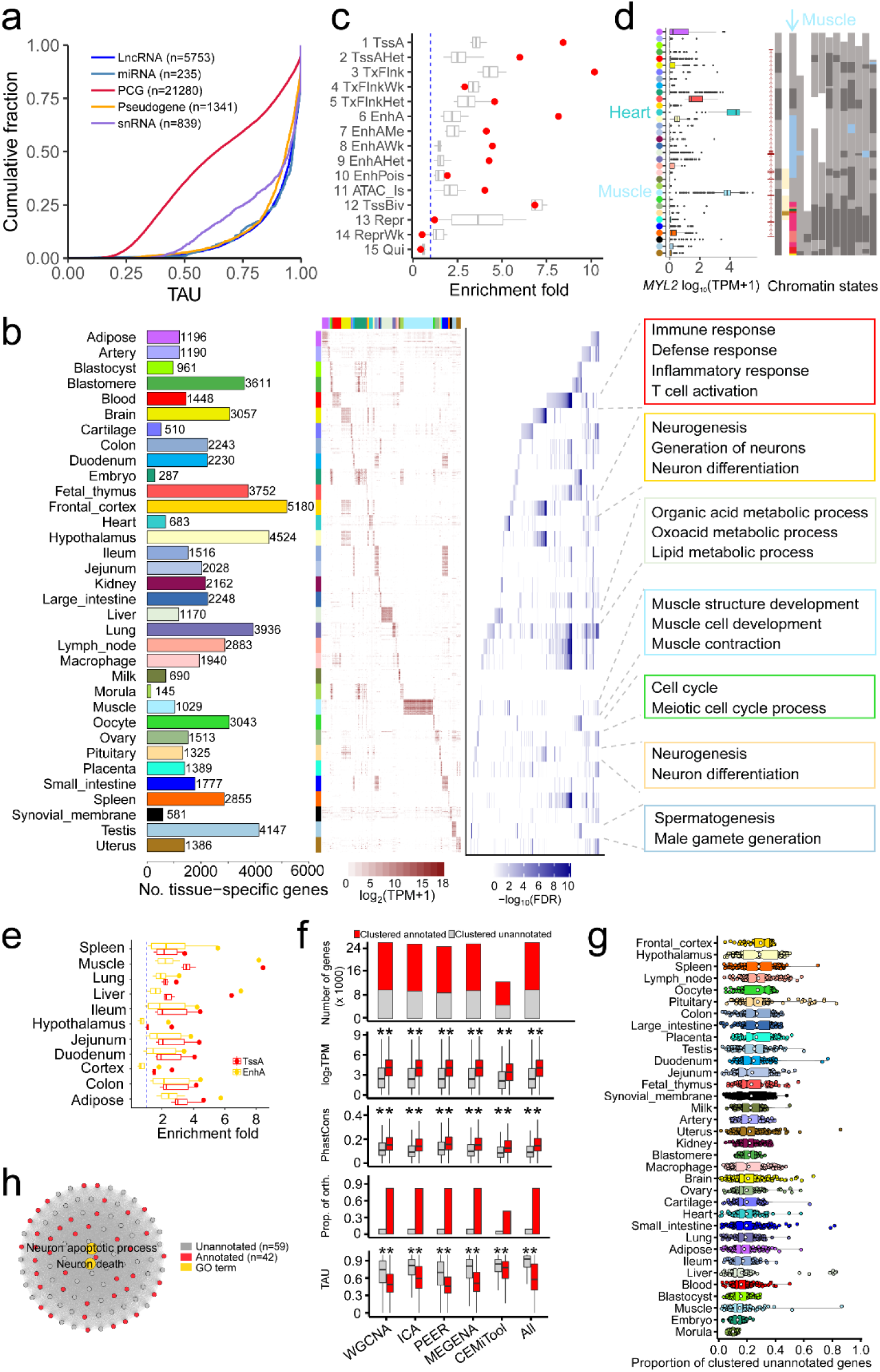
The pig gene expression atlas. (**a**) Tissue-specific expression of five transcript types reflected by the TAU score. PCG: protein coding genes. (**b**) Gene numbers (left), expression pattern (middle, Transcripts Per Million, TPM), and enriched Gene Ontology (GO) terms (right) of tissue-specific genes in 34 tissues. (**c**) Enrichment of muscle-specific genes in 15 chromatin states across 14 pig tissues^17^. The red dots represent chromatin states in muscle. The blue line indicates enrichment fold = 1. (**d**) Expression profiles of *MYL2* gene across 34 tissues (left). The tissue color key is the same as in (**b**). Chromatin state distribution (right) around *MYL2* in 14 pig tissues^17^. In brief, red is for promoter, yellow for enhancers, blue for open chromatin, and grey for repressed regions. (**e**) Enrichment of tissue-specific genes for two active chromatin states across 11 tissues, which have both chromatin states and gene expression data. The dots represent enrichments from matching tissues. TssA is for active TSS (promoter), and EnhA for active enhancers. (**f**) The comparison of genes with and without functional annotation (referred to as “annotated genes” and “unannotated genes”, respectively) in the gene co-expression modules at different biological layers. The gene co-repression analysis was conducted using five complementary methods, including WGCNA, ICA, PEER, MEGENA and CEMiTool. “All” shows the pooled results from the five methods. The functional gene annotation was based on the Gene Ontology database (version 2022-01-18). The plots from top to bottom include gene counts, expression level, PhastCons score from 100 vertebrate genomes, the proportion of orthologous genes in humans, and TAU values. Significant differences between annotated and unannotated genes are obtained using a two-sided *t*-test. ** means *P* < 0.01. (**g**) The proportion of unannotated genes in each gene co-expression modules across 34 tissues. (**h**) An example of gene co-expression module in pituitary, which includes 59 unannotated and 42 annotated genes, respectively. The functional annotated genes are significantly (*P* = 8×10^−3^) enriched in neuron apoptotic processes. The grey edges between genes represent Pearson’s correlations of expression across all 53 samples in pituitary.

**Extended Data Figure 4.**
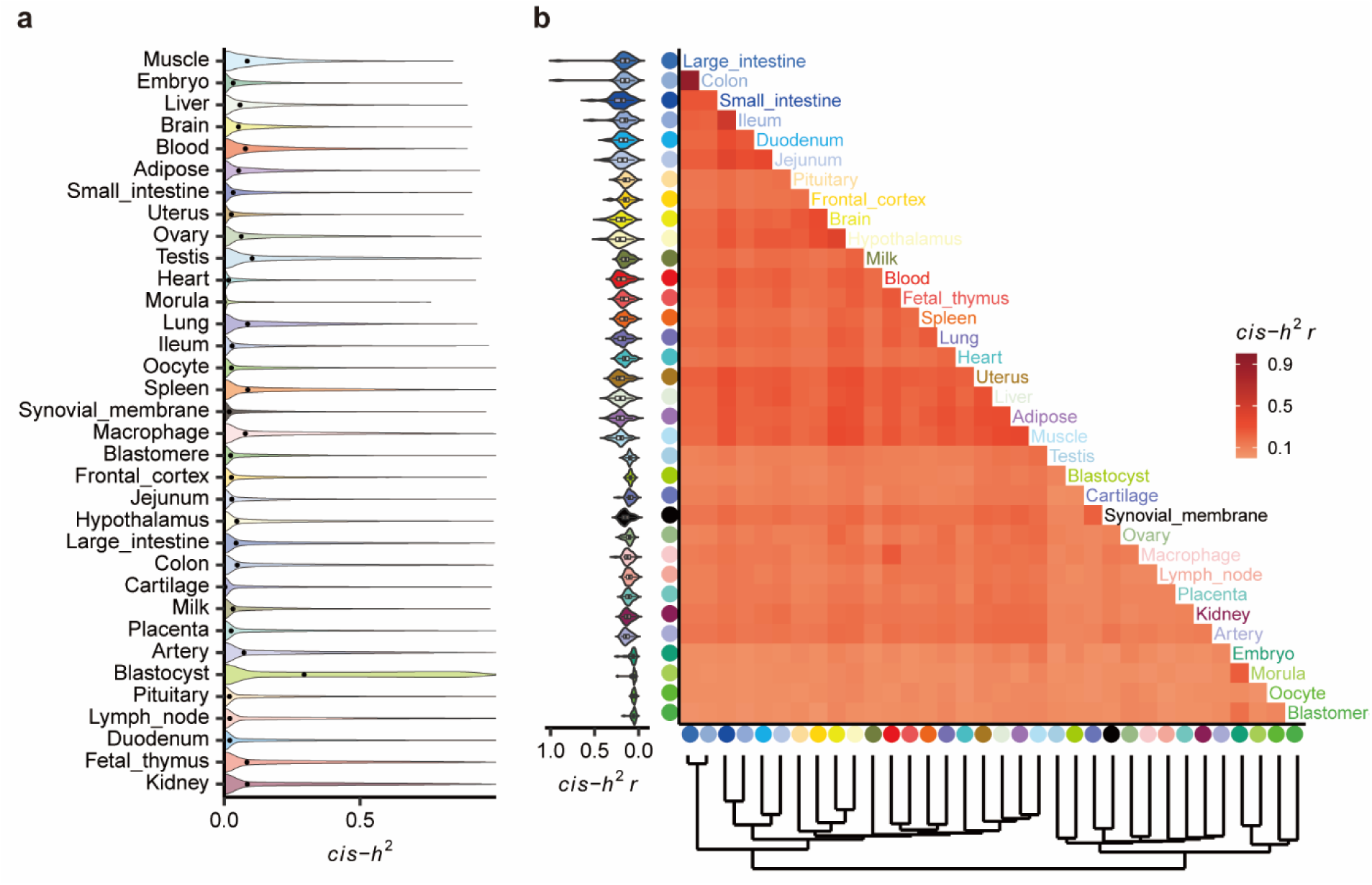
*Cis*-heritability of gene expression across 34 pig tissues. (**a**) Distribution of estimated *cis*-heritability (*cis*-*h*^2^) of gene expression across 34 tissues. The black point represents the median of *cis*-*h*^2^ of all tested genes in a tissue. (**b**) Heatmap of pairwise Pearson’s correlation (*r*) of *cis*-*h*^2^ between tissues. Tissues are grouped by the hierarchical clustering (bottom). The violin plot (left) represents the Pearson’s *r* between the target tissue with the rest.

**Extended Data Figure 5.**
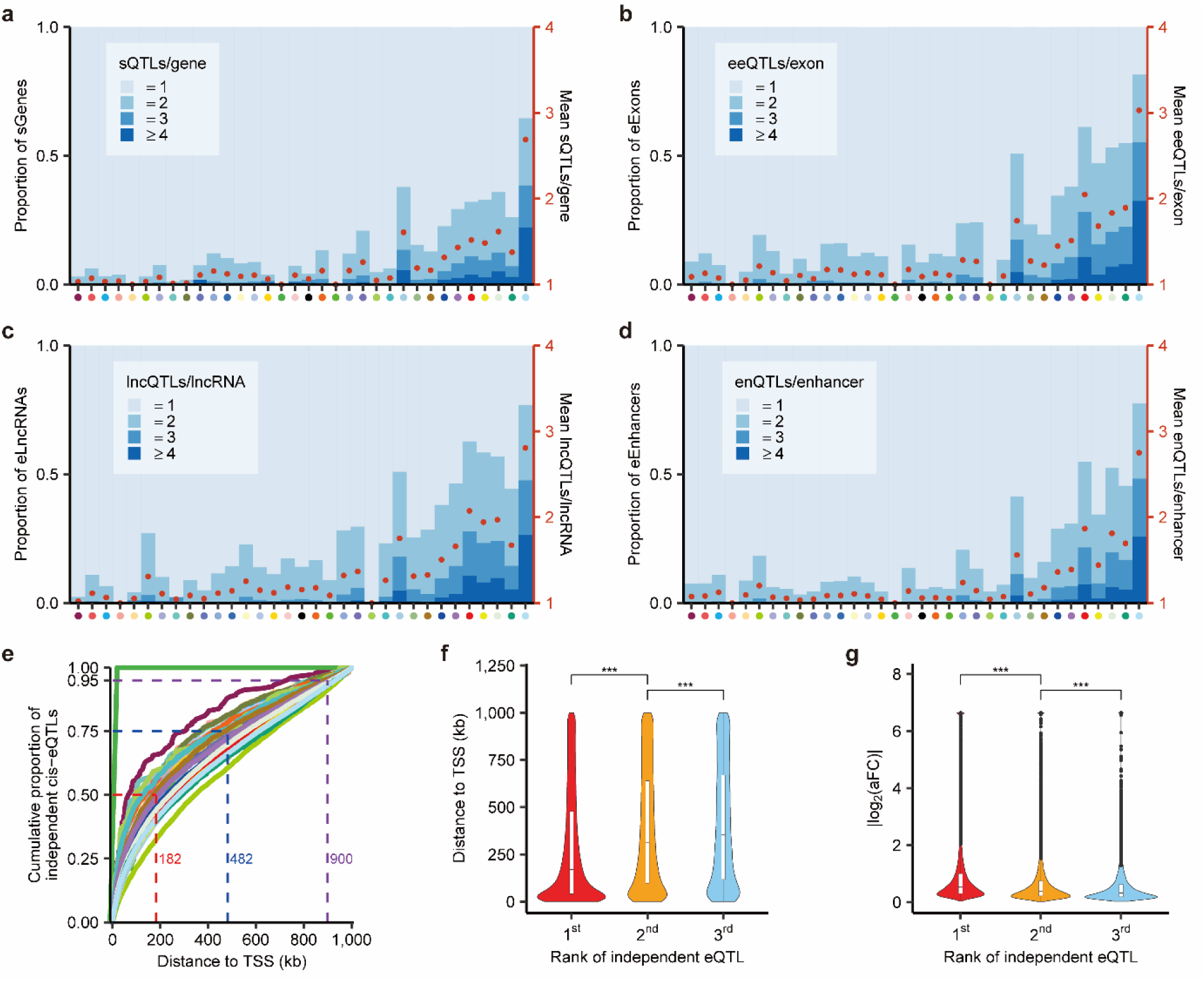
Conditionally independent molecular QTLs (molQTL). (**a-d**) Distribution and average number (red dots, right *y*-axis) of conditionally independent *cis*-sQTL per sGene (**a**), *cis*-eeQTL per eExon (**b**), *cis*-lncQTL per eLncRNA (**c**) and *cis*-enQTL per eEnhancer (**d**) across 34 tissues. The tissues are ordered (from smallest to largest) by sample size. (**e**) Cumulative proportion of distance to transcript start site (TSS) of target genes for conditionally independent *cis*-eQTL in each of 34 tissues. The meaning of color of dots in *x*-axis (**a-d**) and curved lines (**e**) are the same as color key in Fig. 2a. (**f-g**) Distribution of distance to TSS (**f**) and effect size (|log_2_(aFC)|) (**g**) for top three independent *cis*-eQTL per eGene across 34 tissues. The aFC is for allelic fold change. *** represents *P* < 0.001 that obtained from the two-sided Wilcoxon test.

**Extended Data Figure 6.**
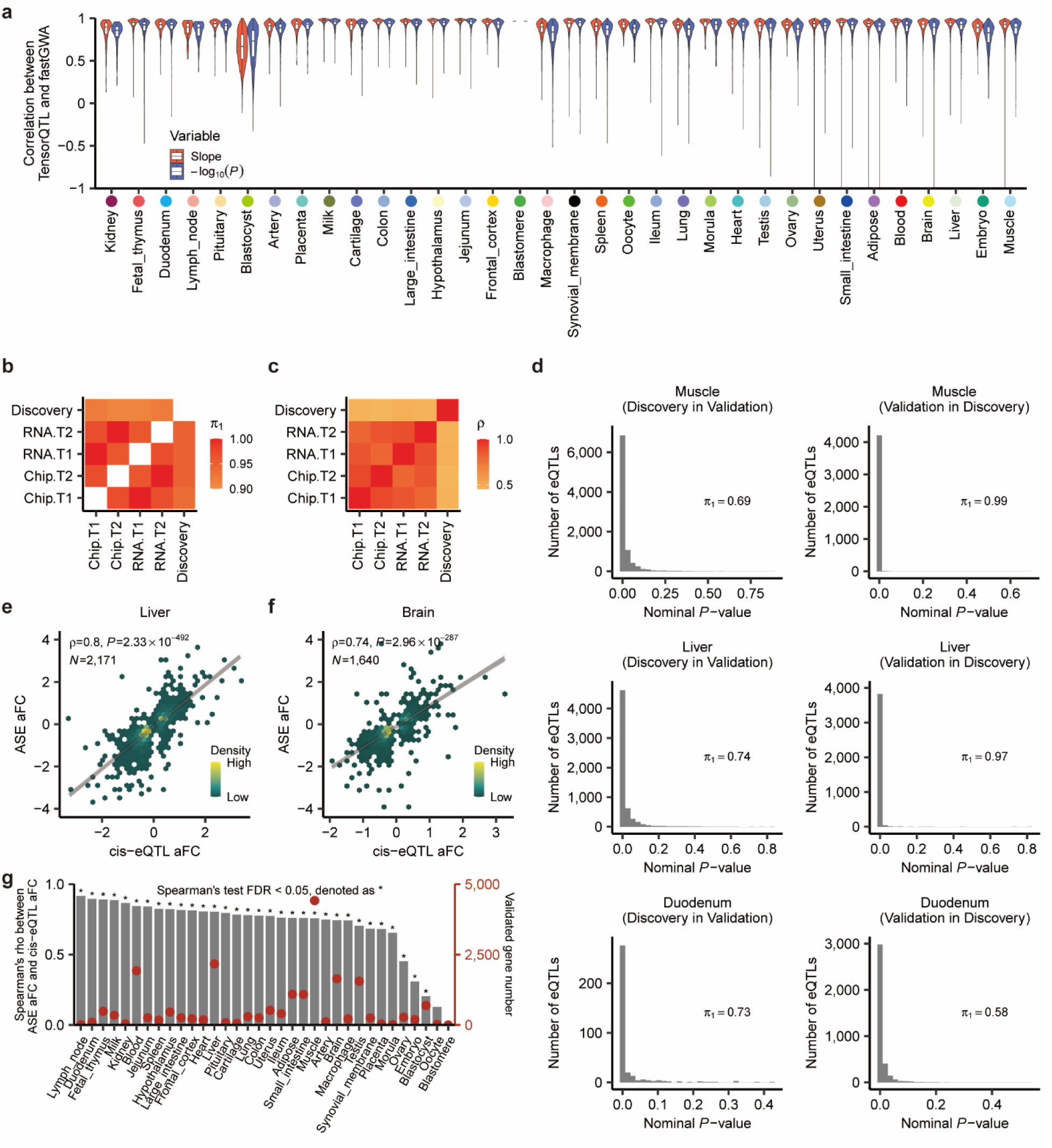
Validation of *cis*-eQTLs. (**a**) Pearson’s correlation (*r*) of *cis*-eQTL summary statistics, including effect size (slope) and significance (-log_10_ adjusted *P*-value), obtained from TensorQTL (v1.0.3) and fastGWA (v1.93.0). TensorQTL implements a standard linear model, while fastGWA implements a mixed linear model considering the kinship among individuals. (**b**) Replication rates (π_1_) of blood *cis*-eQTL between the PigGTEx discovery population (n = 386, Discovery) and the external datasets (n = 179). For π_1_ calculation, rows are discovery populations, and columns are replication populations. The external datasets include whole-blood-cell RNA-Seq data and SNP Chip array (Chip) from 179 animals at two developmental stages (T1 and T2). The prefix “RNA” and “Chip” indicate imputed genotypes from RNA-Seq and SNP array, respectively. (**c**) Spearman’s correlation (ρ) of effect size (z-scores) for blood *cis*-eQTL among the same populations above. (**d**) Replication rates (π_1_) of PigGTEx *cis*-eQTL in external validation datasets of three tissues, including muscle (n_PigGTEx_ = 1,321, n_external_ = 100), liver (n_PigGTEx_ = 501, n_external_ = 100) and duodenum(n_PigGTEx_ = 49, n_external_ = 100). The *x*-axis is the nominal *P*-value of *cis*-eQTL detected from dataset_2_ but is significant in dataset_1_ (i.e., dataset_1_ in dataset_2_). (**e-f**) Spearman’s correlation (ρ) of effect sizes (allelic fold change, aFC) between *cis*-eQTL and matched allele-specific expression (ASE) loci in liver (**e**) and brain (**f**). N indicates number of tested loci. The lines are fitted by a linear regression model using the *geom_smooth* function from ggplot2 (v3.3.2) in R (v4.0.2). (**g**) Spearman’s correlation (ρ) of effect sizes between *cis*-eQTL and matched ASE loci across 34 tissues. Red dots indicate number of tested loci (right *y*-axis).

**Extended Data Figure 7.**
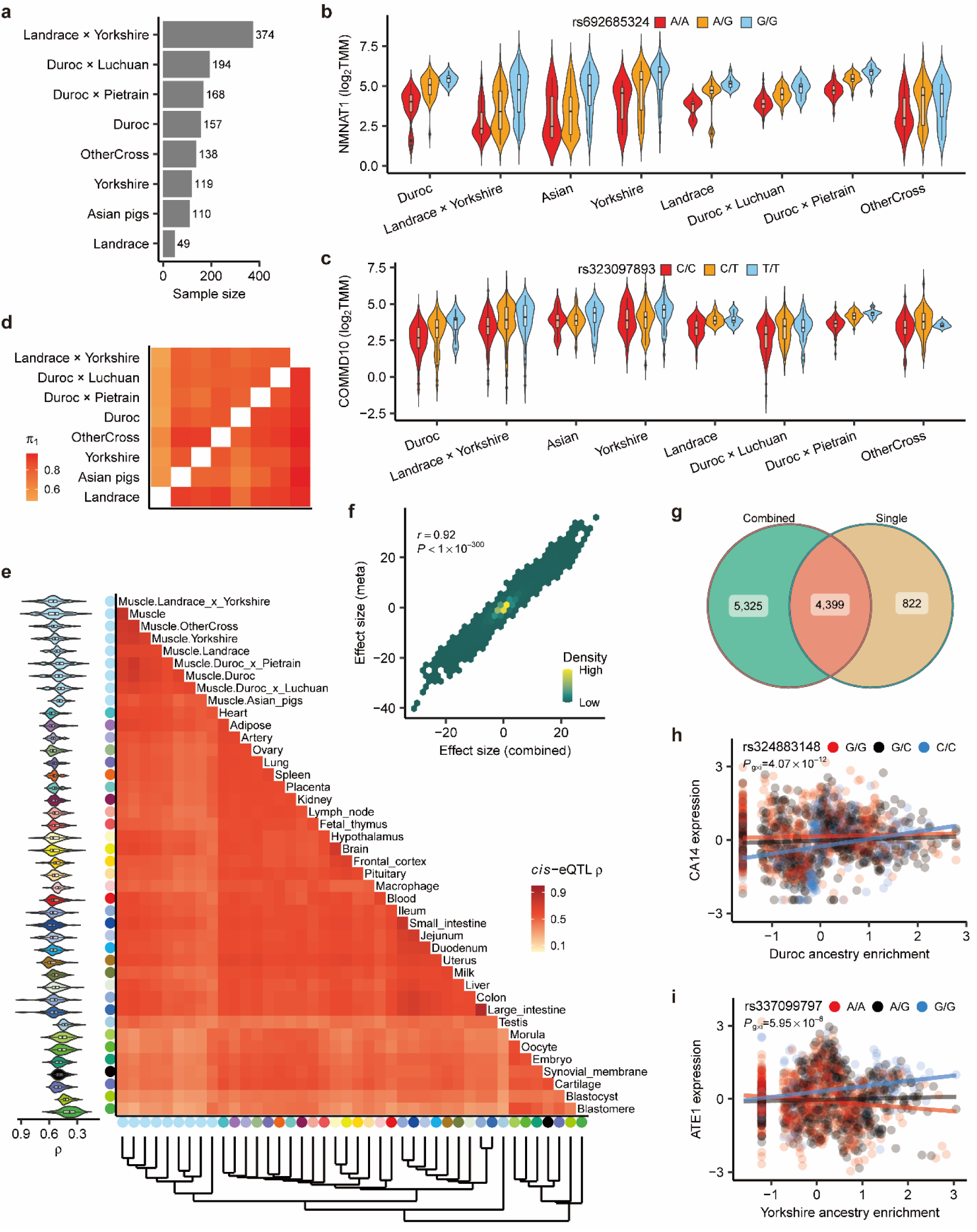
Breed sharing and interaction *cis*-eQTL (bieQTL). (**a**) Sample size of muscle RNA-Seq data across eight breed groups. (**b-c**) Expression levels of *NMNAT1* (**b**) and *COMMD10* (**c**) at three genotypes of *cis*-eQTL in muscle across eight breed groups. (**d**) The *cis*-eQTL discovered in each breed group (rows) that can be replicated (π_1_) across all other breed groups (columns). (**e**) The heatmap of tissues regarding the pairwise Spearman’s correlation (ρ) of *cis*-eQTL effect sizes. Tissues are grouped by the hierarchical clustering (bottom). Violin plot (left) represents the Spearman’s correlation between the target group and the rest. (**f**) Pearson’s correlation (*r*) of effect size between *cis*-eQTL from multi-breed meta-analysis (*y*-axis) and those from the combined muscle population (*x*-axis). (**g**) Overlap of *cis*-eQTL detected from the combined muscle population (Combined) and those detected in single-breed (Single) *cis*-eQTL mapping (shared in at least two breeds). (**h-i**) Examples of bieQTL in muscle. Each dot in (**h**, *CA14*) and (**i**, *ATE1*) represents an individual and is colored by three genotypes. Gene expression levels and ancestry enrichment scores are inverse normal transformed. The lines are fitted by a linear regression model using the *geom_smooth* function from ggplot2 (v3.3.2) in R (v4.0.2).

**Extended Data Figure 8.**
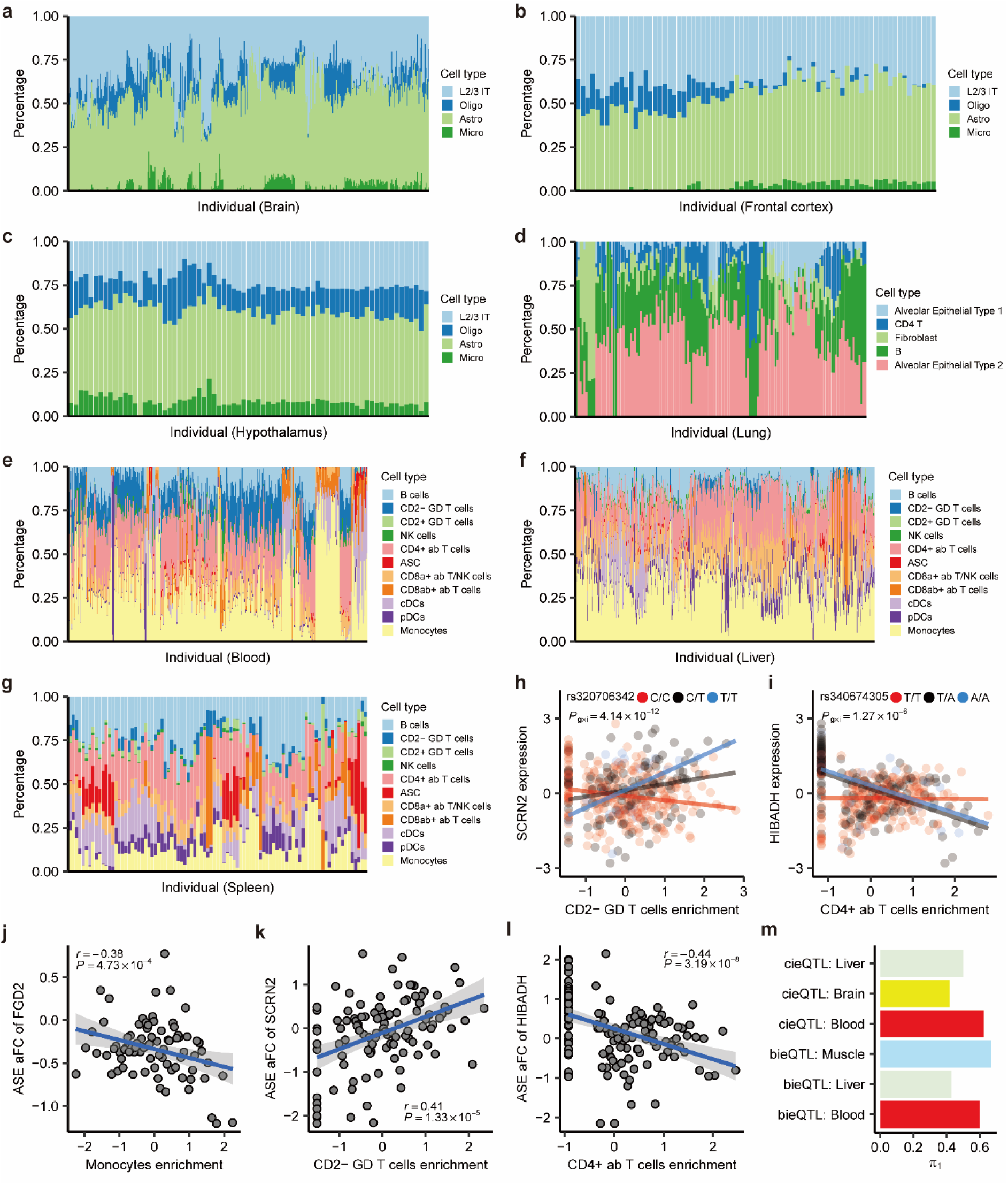
Cell-type enrichment and interaction *cis*-eQTL (cieQTL). (**a-g**) Estimated enrichment (percentage) of major cell types across samples in brain (n = 415) (**a**), frontal cortex (n = 75) (**b**), hypothalamus (n = 73) (**c**), lung (n = 149) (**d**), blood (n = 386) (**e**), liver (n = 501) (**f**), and spleen (n = 91) (**g**). (**h-i**) Examples of cieQTL in blood. Each dot in (**i**, *SCRN2*) and (**j**, *HIBADH*) represents an individual and is colored by three genotypes. Gene expression levels and cell-type enrichment scores are inverse normal transformed. (**j-l**) Pearson’s correlation (*r*) between allele-specific expression (ASE) effect sizes (allelic fold change, aFC) and specific cell-type enrichment scores for *FGD2* with monocytes (**j**), *SCRN2* with CD2-GD T cells (**k**) and *HIBADH* with CD4+ ab T cells in blood (**l**). The lines are fitted by a linear regression model using the *geom_smooth* function from ggplot2 (v3.3.2) in R (v4.0.2). (**m**) ASE validation rate (π_1_) of breed/cell-type interaction QTL (bieQTL and cieQTL) across tissues with ≥ 5 detectable bieQTL or cieQTL.

**Extended Data Figure 9.**
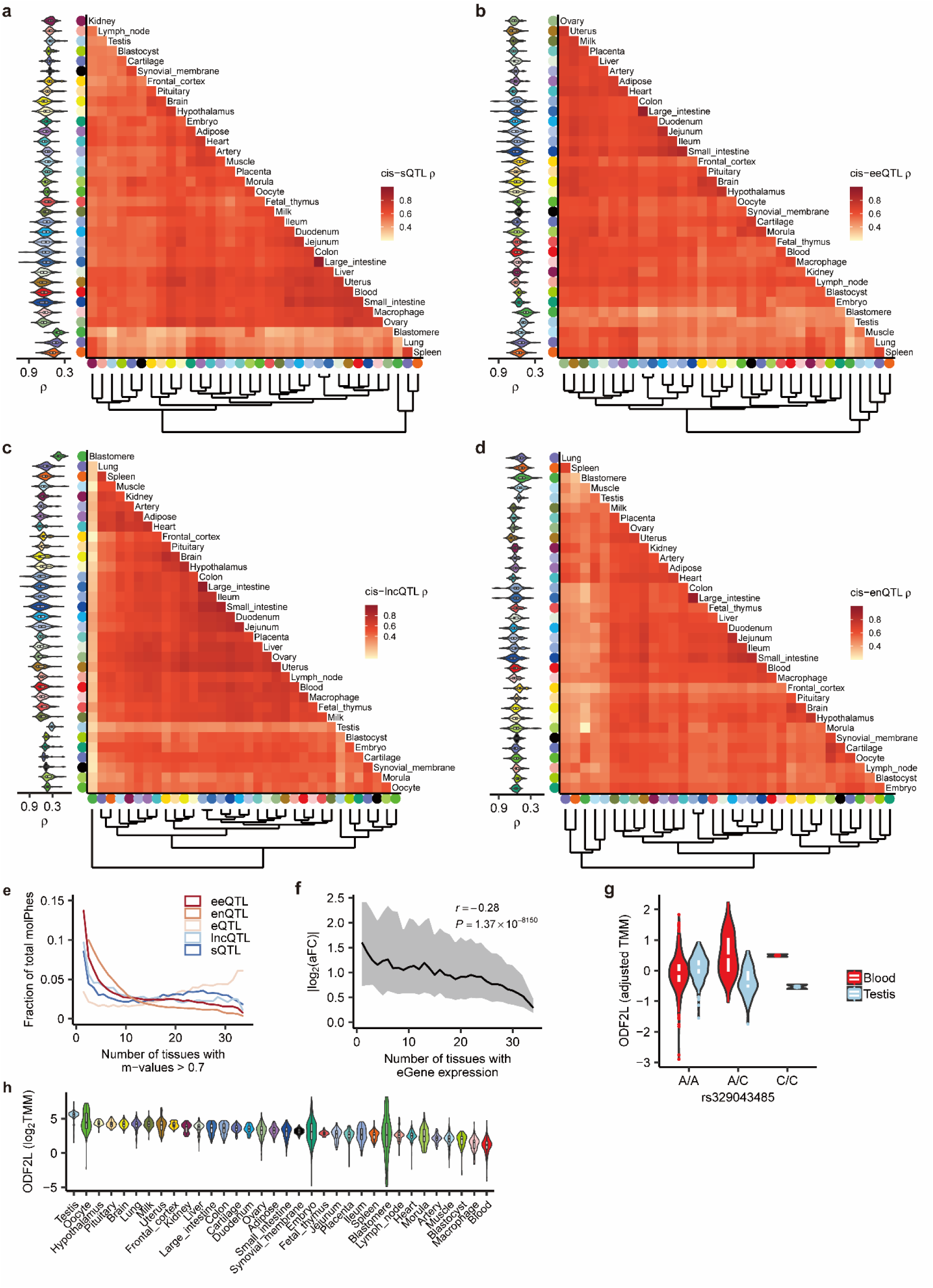
Tissue sharing and specificity patterns of molecular QTL (molQTL). (**a-d**) The heatmap of tissues regarding the pairwise Spearman’s correlation (ρ) of molQTL effect sizes, i.e., *cis*-sQTL (**a**), *cis*-eeQTL (**b**), *cis*-lncQTL (**c**), and *cis*-enQTL (**d**). Tissues are grouped by the hierarchical clustering (bottom). Violin plot (left) represents the Spearman’s correlation between a tissue and other tissues. (**e**) Distribution of number of tissues having METASOFT activity (M-value > 0.7) for each of molQTL. MolPhe: molecular phenotypes. (**f**) Pearson’s correlation (*r*) between number of tissues an eGene expressed in (Transcript Per Million, TPM > 0.1) and its *cis*-eQTL effect sizes (|aFC(log_2_)|). The aFC is for allelic fold change. The line and shading indicate the median and interquartile range, respectively. (**g**) Expression levels (adjusted TMM) of *ODF2L* at three genotypes of top *cis*-eQTL (rs329043485) in blood and testis. TMM: Trimmed Mean of M-value normalized expression levels. There are 337, 47 and 2 samples for A/A, A/C and C/C genotypes in blood, respectively, and 148, 34 and 2 in testis. (**h**) Expression levels (log_2_TMM) of *ODF2L* across 34 tissues. Tissues are ordered (from smallest to largest) by the median expression values.

**Extended Data Figure 10.**
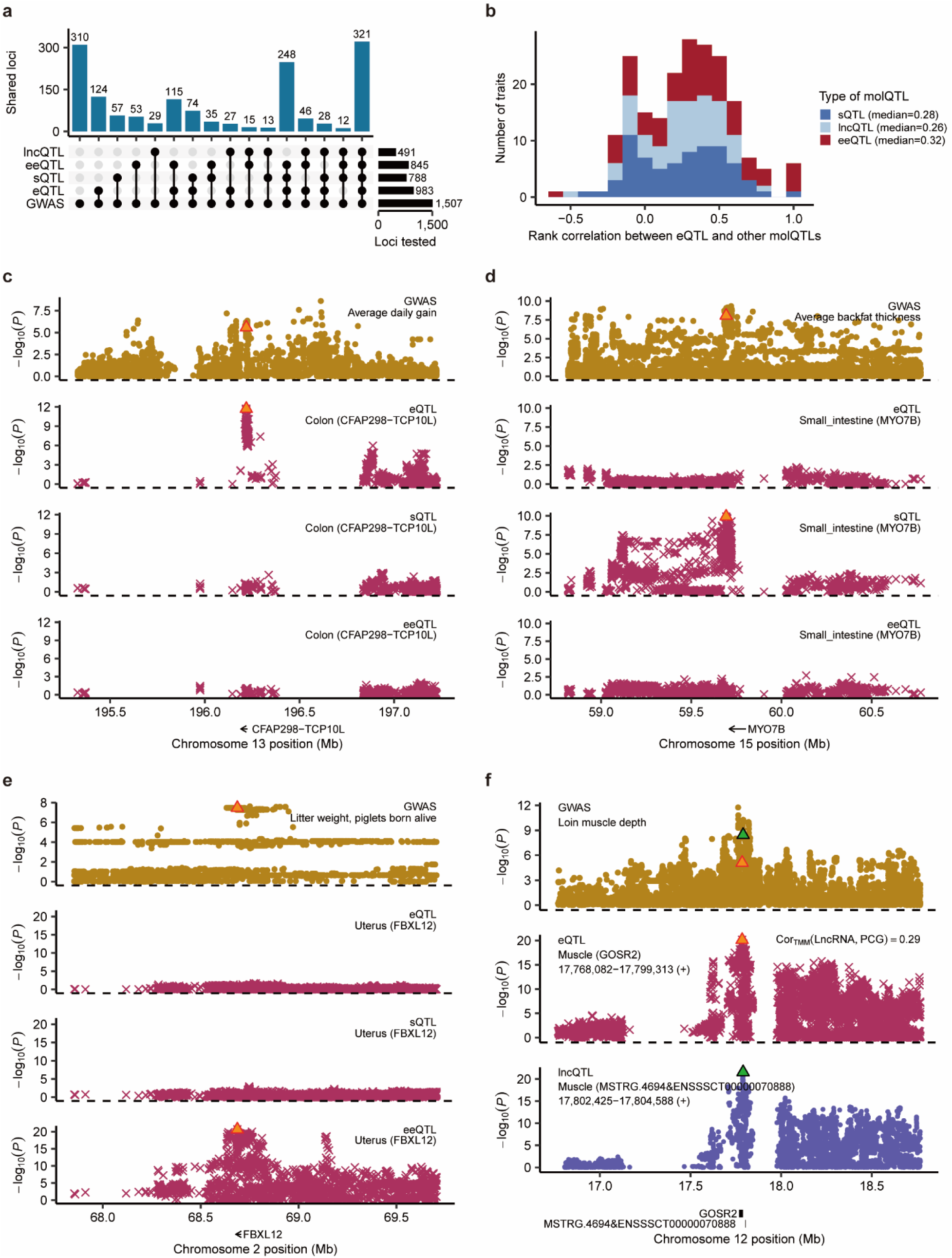
Complementarity of molecular QTL (molQTL) in interpretating GWAS loci. (**a**) Number of GWAS loci linked to *cis*-eQTL, *cis*-sQTL, *cis*-eeQTL, and *cis*-lncQTL in 34 tissues based on four different integrative methods, including colocalization (fastEnloc), Mendelian randomization (SMR), single-tissue transcriptome-wide association studies (TWAS, S-PrediXcan) and multi-tissue TWAS (S-MultiXcan). (**b**) Distribution of rank correlations between tissue-relevance-scores derived from *cis*-eQTL and those from *cis*-sQTL, *cis*-lncQTL and *cis*-eeQTL across 86 GWAS traits with significant colocalizations for at least one molecular phenotype. (**c**) Significant SMR signals (*P*_SMR_ = 9.16×10^−5^, *P*_HEIDI_ = 0.9) between GWAS loci of average daily gain (ADG) and *cis*-eQTL of *CFAP298-TCP10L* in colon on chromosome 13, but not for its *cis*-sQTL or *cis*-eeQTL. The orange triangle represents the top *cis*-eQTL of *CFAP298-TCP10L* in colon. (**d**) Significant SMR signals (*P*_SMR_ = 1.78×10^−5^, *P*_HEIDI_ = 0.07) between GWAS loci of average backfat thickness (BFT) and *cis*-sQTL of *MYO7B* in small intestine on chromosome 15, but not for its *cis*-eQTL or *cis*-eeQTL. (**e**) Significant SMR signals (*P*_SMR_ = 1.78×10^−6^, *P*_HEIDI_ = 0.97) between GWAS loci of litter weight (LW, piglets born alive) and *cis*-eeQTL of *FBXL12* in uterus on chromosome 2, but not for its *cis*-eQTL or *cis*-sQTL. (**f**) Significant SMR signals (*P*_SMR(lncQTL-GWAS)_ = 4.49×10^−7^, *P*_SMR(eQTL-GWAS)_ = 5.45×10^−5^, *P*_SMR(lncQTL-eQTL)_ = 4.62×10^−7^) among GWAS loci of loin muscle depth (LMD), *cis*-lncQTL of *MSTRG.4694&ENSSSCT00000070888*, and *cis*-eQTL of *GOSR2* in muscle on chromosome 12. *MSTRG.4694&ENSSSCT00000070888* is a lncRNA gene located on the 3112 bp downstream of *GOSR2*, where the Pearson’s correlation of their normalized expression levels (Trimmed Mean of M-value, TMM) is 0.29 in muscle. The orange and green triangles in the top GWAS Manhattan plot represent the top molQTL of *GOSR2* and *MSTRG.4694&ENSSSCT00000070888*, respectively.

## Online Methods

### Ethics

Not applicable because no biological samples were collected and no animal handling was performed for this study.

### RNA-Seq data analysis and molecular phenotype quantification

In total, we gathered 11,323 publicly accessible raw RNA-Seq datasets (fastq files), representing 9,530 distinct samples (downloaded from NCBI SRA database by February 26^th^, 2021), of which 98.13% were generated using the Illumina platform. We removed 121 embargoed RNA-Seq samples, and then processed all the remaining RNA-Seq samples using a uniform pipeline. Briefly, we first trimmed adaptors and discarded reads with poor quality using Trimmomatic (v0.39) with the following parameters: TruSeq3-PE.fa:2:30:10, LEADING:3, TRAILING:3, SLIDINGWINDOW:4:15 and MINLEN:36 ^45^. We then aligned clean reads to the Sscrofa11.1 (v100) pig reference genome using STAR (v2.7.0) with the following options: outFilterMismatchNmax 3, outFilterMultimapNmax 10 and outFilterScoreMinOverLread 0.66 ^46^. We kept 8,262 samples with more than 500K clean reads and uniquely mapping rates ≥ 60% for subsequent analysis (Table S1). We extracted the raw read counts of 31,871 Ensembl (Sscrofa11.1 v100) genes by featureCounts (v1.5.2)^47^ and obtained their normalized expression (i.e., TPM) using Stringtie (v2.1.1)^48^. We removed 544 samples in which less than 20% of all annotated genes were expressed (TPM ≥ 0.1), resulting in 7,597 samples. We then visualized the variance in gene expression among samples using t-distributed stochastic neighbor embedding (*t*-SNE)^49^ with distance = (1-*r*), where *r* is Pearson’s correlation coefficient of gene expression. After filtering out outliers within each of the tissues, we eventually kept 7,095 samples for subsequent analysis (Table S1). We employed MEGA X^50^ to build a Neighbor-joining tree of these samples based on TPM and then visualized it by iTOL (v6)^51^.

For PCG expression, we considered 21,280 PCGs from the Ensembl annotation of the porcine genome (Sscrofa11.1 v100). For exon expression of PCGs, we extracted raw read counts of 290,536 exons by featureCounts (v1.5.2)^47^ and then normalized them as TPM. To explore enhancer expression, we downloaded the previously predicted enhancers (strong active enhancers, EnhA) from 14 pig tissues^17^. We merged these enhancer regions across tissues using bedtools (v2.30.0)^52^, resulting in 158,998 non-redundant enhancer regions. We obtained raw read counts of these non-redundant enhancer regions from all 7,095 RNA-Seq samples by featureCounts (v1.5.2)^47^, followed by TPM normalization. For lncRNA expression, we obtained 17,162 lncRNAs predicted from 33 Iso-Seq datasets, representing 10 tissues from 4 animals by using FEELnc^53^. We applied the same approach to extract and normalize lncRNA expression as above.

For alternative splicing, we utilized Leafcutter (v0.2.9)^54^ to quantify excision levels of introns and then to identify splicing events within each tissue as described in the following: (1) Converting aligned bam files from STAR (v2.7.0) to junction files using the script *bam2junc.sh*; (2) Generating intron clusters using the script *leafcutter_cluster.py* with parameters: --min_clu_reads 30, -- min_clu_ratio 0.001, --max_intron_len 500000, --num_pcs 15, and then mapping them to genes by the *map_clusters_to_genes.R* script with exon coordinates extracted from the Ensembl annotation file (v100); (3) Discarding introns without any read count in more than 50% of samples or with fewer than *max*(10, 0.1n) unique values, where n is the sample size; (4) Filtering out introns with low complexity : ∑*_i_*(|z*_i_*| < 0.25) ≥ n-3 and ∑*_i_*(|z*_i_*| > 6) ≤ 3, where z*_i_* is the z-score of the *i*th cluster read fraction across individuals (*_perind.counts.gz files); (5) Using *prepare_phenotype_table.py* script to normalize filtered counts and convert them to BED format, where start/end positions correspond to the TSS of corresponding genes. Furthermore, we normalized excision levels of introns as percent splicing (PSI) values.

### Tissue-specific gene expression and co-expression analysis

To explore the tissue specificity of gene expression, we only considered 34 tissues with over 40 RNA-Seq samples. We calculated eight distinct metrics to quantify the tissue specificity of gene expression, including Counts, Gini, JS, Roku, Shannon, Simpson, Spm and TAU (τ)^55^. To identify genes with specifically high expression in a tissue, we conducted differential gene expression analysis by comparing the target tissue with the other tissues using the limma (v3.51.2) R package^56^. We considered genes with LogFC > 2 and FDR < 0.05 as differentially expressed between tissues. To validate the tissue-specific genes, we downloaded 15 chromatin states predicted in 14 major pig tissues^17^. We conducted the chromatin state enrichment analysis of tissue-specific genes as described in ChromHMM (v1.22)^57^: (C/A)/(B/D), where A is the number of bases in the chromatin state, B is the number of bases in tissue-specific genes, C is the number of bases in both the chromatin state and tissue-specific genes, and D is the number of bases in the entire genome. We calculated the statistical significance of enrichment using Fisher’s exact test. We performed gene co-expression analysis within each of 34 tissues using five complementary methods, including WGCNA (v1.69)^58^, ICA (v1.0.2)^59^, PEER (v1.3)^60^, MEGENA (v1.3.7)^61^, and CEMiTool (v1.8.3)^62^ with default parameters. We adjusted gene expression for hidden confounding factors using PEER factors and genotype PCs (see below in molQTL mapping section) and then used the residuals to infer gene co-expression modules by each of the five methods^63^. We defined whether a gene is annotated in the GO database or not using the *getBM* function in biomaRt R package (v2.48.0)^64^. We employed clusterProfiler (v4.0) to conduct the gene functional enrichment analysis based on the GO database^65^. We visualized the gene co-expression network using Gephi (v0.9.2)^66^.

### Bioinformatics analysis of WGBS data

We downloaded 245 publicly available WGBS data (fastq files) from NCBI SRA and CNCB GSA by July 11^th^, 2021, representing 29 tissues, all generated using Illumina sequencers. We processed all these data using a uniform pipeline as described in the following. We first used FastQC (v0.11.9) (https://www.bioinformatics.babraham.ac.uk/projects/fastqc/) to evaluate the read quality and removed reads with low quality using Trim Galore (v0.4.5) (https://www.bioinformatics.babraham.ac.uk/projects/trim_galore/) with parameters: --max_n 15 -- quality 20 --length 20 -e 0.1. We then aligned clean reads to the pig reference genome (Sscrofa11.1.100) using Bismark (v0.19.0) with default parameters and removed PCR duplicates from the mapped reads using the *deduplicate_bismark* function^67^. We obtained the methylation level of cytosines using the *bismark_methylation_extractor* (--ignore_r2 6) routine and removed CpG sites represented by less than five reads. Ultimately, we retained 182 samples with more than 10 million clean reads and unique mapping rate > 40%, representing 10 tissues and cell lines. We then removed data from corpus luteum as it had less than five samples. We visualized the variance in DNA methylation among samples using *t*-SNE, and removed samples from two cell lines, fibroblast and iPSC, as they were not clustered together or had no corresponding cell types in the current RNA-Seq dataset. Finally, we kept 166 samples from seven tissues to detect tissue-specific hypomethylated regions (HMR) and allele-specific methylation (ASM) loci. To detect tissue-specific HMR, we used SMART2 (v2.2.8)^68^ with options: -t DeNovoDMR -MR 0.5 -AG 1.0 -MS 0.5 -ED 0.2 -SM 0.6 -CD 500 -CN 5 -SL 20 -PD 0.05 -PM 0.05 -AD 0.3. For ASM analysis, we employed Methpipe (v4.1.1)^69^. We first calculated the ASM score of each CpG site using the *allelicmeth* function and then obtained significant (FDR < 0.01) ASMs using the *amrfinder* function.

### Bioinformatics analysis of Hi-C data

We downloaded Hi-C data of five samples from five different pig tissues (i.e., adipocyte, ear, embryo, liver and muscle) from NCBI SRA to identify TAD and Hi-C contacts (Table S8). After trimming adapter sequences and low-quality reads using Trim Galore (v0.6.7) (https://www.bioinformatics.babraham.ac.uk/projects/trim_galore/) with parameters: --q 20 --paired -- max_n 15 --clip_R2 3, we aligned clean reads to the pig reference genome (Sscrofa11.1.100) using BWA (v0.7.17) with default settings^70^. We built Hi-C contact matrices at 500 kb resolution with Juicer (v1.6)^71^ and identified TAD with Arrowhead (v1.22.01)^71^. We only considered TADs with FDR < 0.01 for the downstream analysis. We applied pyGenomeTracks (v3.6)^72^ to visualize gene examples after converting .hic files to .cool files and then to .h5 files using hicConvertFormat (v3.7.1)^73^.

### Single-cell RNA-Seq and cell-type deconvolution analysis

We obtained 13 raw single-cell RNA-Seq data of five brain and eight lung samples from the CNGB Sequence Archive (CNSA) of the China National GeneBank DataBase (CNGBdb) under accessions: CNP0000686 and CNP0001486, respectively. We also obtained the processed single-cell RNA-Seq data of seven PBMC samples from USDA Ag Data Commons^74^.

### Identification of cell types

For five brain regions (frontal lobe: FL, parietal lobe: PL, temporal lobe: TL, occipital lobe: OL, and hypothalamus: HT), we used the Cell Ranger Single Cell Software Suite (v3.0.2) (https://support.10xgenomics.com/single-cell-gene-expression/software/pipelines/latest/what-is-cell-ranger) to process and analyze the raw 10x Genomics sequencing data. We used the *cellranger mkgtf* tool to keep all gene types in the gene annotation .gtf file (Ensembl v100) and the *cellranger mkref* tool to build a pig custom reference using both the generated GTF and Sscrofa11.1 (v100) genome assembly. We generated the raw count matrices using the *cellranger count* tool and then performed all subsequent analyses using the Seurat (v3.2.3) R package^75^. Overall, we obtained 2,465, 1,312, 4,974, 3,756, and 1,462 cells in FL, HT, OL, PL and TL, respectively, with the following criteria: 1) cells with gene counts over 200; and 2) cells with mitochondrial counts less than 5% of the total counts. We calculated gene expression using the *LogNormalize* method, implemented in the *Normalization* function and selected 2,000 genes as highly variable genes (HVG) using the *FindVariableFeatures* function with default parameters. Using the *FindIntegrationAnchors* and *IntegrateData* functions (dim = 1:20), we then integrated the five gene count matrices of HVG into one *Seurat* object, while correcting for batch effects. We applied the *ScaleData* function to scale the integrated data and then used it for principal components analysis (PCA) using the *RunPCA* function. Based on the elbow plot generated by the *JackStraw* function, we used the first 20 PCs for cell clustering and Uniform Manifold Approximation and Projection (UMAP) analysis with the *RunUMAP* function. We constructed a shared nearest neighbor (SNN) graph of cells with the *FindNeighbors* function and determined cell clusters with *FindClusters* at a resolution of 0.4. We then applied the UMAP algorithm to visualize the cell clustering. We utilized Azimuth (v0.4.0)^76^ with a human motor cortex reference to label the cell clusters and assigned all cells to eight known brain cell types.

For lung tissues, we employed the DNBelab C Series scRNA analysis software to process the raw sequencing data generated from the DNBelab C4 sequencing platform^77^, including creating a reference genome database and producing the gene expression matrix. We then applied the same pipeline and standards as described above to perform the following single-cell analysis. Collectively, 1,744, 966, 240, 1,317, 1,749, 10,000, 10,000, and 1,596 cells passed the quality control across eight samples. We assigned 35 cell types according to the human lung reference v1^78^, implemented in Azimuth (v0.4.0)^76^. For PBMC, we directly used cell clusters provided by Herrera-Uribe et al.^74^ for subsequent analysis.

### Cell type deconvolution Analysis

We selected four, five, and ten major cell types from brain, lung, and PBMC, respectively, and then applied CIBERSORTx^79^ to estimate the fraction of these cell types in bulk RNA-Seq samples from seven tissues, including brain, frontal cortex, hypothalamus, lung, blood, liver, and spleen. We extracted 150 cells from each cell cluster using the *subset* function, implemented in Seurat (v3.0.2)^75^, to create a signature matrix using the CIBERSORTx online tool by the *custom* option with default parameters. We then uploaded the gene expression (TPM) matrix of bulk RNA-Seq samples as the mixture file. We imputed cell fractions based on the signature and mixture files by running the *Impute Cell Fractions* analysis with the *custom* mode and used the permutation test (100 times) to determine the significance level.

### Genotype imputation reference panel

To develop a genotype imputation reference panel for imputing sequence variant genotypes into the RNA-Seq samples, we downloaded 1,307 public WGS samples from NCBI SRA by March 18^th^, 2021, and newly generated an additional 510 WGS samples (Table S2), representing five main pig populations worldwide, i.e., Suidae but not Sus scrofa (SUI, n = 45), European wild boar (EUW, n = 54), European domestic pig (EUD, n = 855), Asian wild boar (ASW, n = 80), and Asian domestic pig (ASD, n = 783). We processed and analyzed the WGS data using a uniform pipeline as described in the following. Briefly, we filtered the raw sequence reads by Trimmomatic (v0.39)^45^, and then mapped clean reads to Sscrofa11.1 using BWA-MEM (v0.7.5a-r405) with default parameters^70^. We marked duplicated reads by Picard (v2.21.2) (http://broadinstitute.github.io/picard/). We removed 213 samples with low read depth (< 5x), one sample with low genome coverage (< 75%), and one sample with incomplete data. Finally, we kept 1,602 samples for jointly calling variants using the Genome Analysis Toolkit (GATK) (v4.1.4.1)^80^ with parameters: QD> 2, MQ < 40, FS > 60, SOR > 3, MQRankSum < −12.5 and ReadPosRankSum < −8, yielding ~214 million SNPs. We removed SNPs with MAF < 0.01 and/or missing rate > 0.9 using bcftools (v1.9)^81^ and employed Beagle 5.1 to phase the filtered variants and impute sporadically missing genotypes^82^. Finally, a total of 42,523,218 SNPs were retained in the current version of PGRP (v1).

### Genotyping and imputation of RNA-Seq samples

We employed the GATK (v4.0.8.1) to call SNPs at known loci in the dbSNP database (build 150), from 7,095 RNA-Seq samples, according to the recommended settings of the best practice guidelines^80^. We filtered out low-quality SNPs using the filtering option: FS > 30.0 & QD < 2.0 & DP > 3.0. We then imputed the filtered SNPs on autosomes to sequence level with Beagle (v5.1)^82^ using haplotypes from the PGRP as a reference. Finally, we obtained 7,008 samples that were genotyped and imputed successfully. We filtered out variants with MAF < 0.05 and model-based imputation accuracy (DR^2^) < 0.85, resulting in 3,087,268 SNPs for the molQTL mapping. We obtained 12,207 linkage disequilibrium (LD) independent SNPs using PLINK (v1.90)^83^ with parameters: --indep pairwise 1000 5 0.2 ^83^, and then conducted a principal components (PC) analysis for all 7,008 samples using these LD-independent SNPs.

To evaluate the accuracy of genotype imputation from RNA-Seq, we first examined 36 samples with both RNA-Seq and WGS data from four tissues of two crossbred populations, i.e., Landrace×Yorkshire (n = 25) and Duroc×Diannanxiaoer (n = 11). We calculated accuracy of imputation as the concordance between genotypes imputed from RNA-Seq and those directly called from WGS ^84^. In addition, we collected 37 Duroc and 179 Duroc×Landrace×Yorkshire pigs, which had both RNA-Seq and 50k SNP array. We imputed genotypes from 50k SNP array only using the same pipeline as described above and then compared them with those imputed from RNA-Seq only by calculating the concordance rate.

### Breed prediction for RNA-Seq samples

There were 3,684 public RNA-Seq samples without breed information. We thus predicted their breed composition using the imputed SNPs *via* the pipeline summarized in Extended Data Fig. 1i. In brief, we estimated the breed composition of all 7,008 RNA-Seq samples based on 5,000 randomly selected LD-independent SNPs using ADMIXTURE (v1.3.0)^85^. According to the estimated ancestry proportion of samples with known breed information from three pure breeds (Duroc, Landrace, and Yorkshire) and two crossbred populations (Landrace×Yorkshire and Duroc×Landrace×Yorkshire) at different K values in the admixture analysis, we decided to choose K = 5 for further analysis. We identified samples with more than 10% ancestry proportion from ≥ 2 ancestries as crossbreeds, while the remaining samples were considered as pure breeds. For crossbreeds and other pure breeds, we trained a machine learning model (implemented in RandomForest R package^86^) using samples with known breed labels to predict their breed information. We repeated the three-fold cross-validation 10 times to evaluate the prediction accuracy.

### Molecular QTL (molQTL) mapping

To gain a comprehensive understanding of the genetic regulatory mechanism underpinning molecular phenotypes in pigs, we performed molQTL mapping for five molecular phenotypes in each of 34 tissues, including PCG expression, exon expression, lncRNA expression, enhancer expression and alternative splicing (PSI). In addition, we exploited interactions of genotypes with cell type and breed compositions.

### Detection of duplicated RNA-Seq samples within each tissue

To remove samples from the same individuals within each tissue, we first calculated the identity-by-state (IBS) distance among samples from LD-independent SNPs using PLINK (v1.90). IBS = (IBS2 + 0.5*IBS1) / (IBS0 + IBS1 + IBS2), where IBS0 is the number of non-missing variants with IBS = 0 (two different alleles), IBS1 is the number of non-missing variants with IBS = 1 (one shared allele), and IBS2 is the number of non-missing variants with IBS = 2 (two shared alleles). We set an IBS distance cutoff of 0.9 to deem samples as duplicates and kept the one with the largest number of expressed genes for subsequent analysis. We then removed an average of 54 duplicated samples (from 1 in kidney to 455 in blood) within each tissue, resulting in 5,457 samples in 34 tissues. To validate whether duplicate samples could be identified correctly, we calculated IBS distance of 25 RNA-Seq samples from 9 individuals. We observed that samples from the same individual could be grouped together well according to IBS cutoff of 0.9. In addition, we calculated the Pearson’s correlation of IBS calculated from imputed genotypes and those from WGS or SNP array from the same individuals.

### Covariates for molQTL mapping

For molQTL mapping within each of the 34 tissues, we only considered SNPs with MAF ≥ 5% and minor allele count (MAC) ≥ 6, resulting in an average of 2,705,637 SNPs (ranging from 1,815,729 in synovial membrane to 3,004,852 in muscle). We then computed genotype PCs based on the filtered SNPs within each of the tissues using SNPRelate (v1.26.0)^87^. We used the top five and ten genotype PCs to account for the population structure among samples in tissues with < 200 and ≥ 200 samples, respectively (Extended Data Fig. 2f). To account for technical confounders among RNA-Seq samples (e.g., hidden batch effects and other technical or biological factors), we used the Probabilistic Estimation of Expression Residuals (PEER) method, implemented in peer R package^60^, to estimate a set of latent covariates within each of the 34 tissues based on gene expression matrices. We obtained a total of 60 PEER factors in each tissue and assessed their relative contributions (i.e., factor weight variance) to gene expression variation using the *PEER_getAlpha* function. As in the human GTEx^7^, we decided to use the top ten PEER factors for each tissue as covariates when conducting molQTL mapping for PGC, exon, lncRNA and enhancer expression. For *cis*-sQTL mapping, we estimated and fitted ten PEER factors from the splicing quantifications (i.e., PSI) of genes within each tissue. To understand whether known covariates can be captured by PEER factors, we fitted a linear regression model to estimate the proportion of variance in known confounders (e.g., data source, sequencing platform, and estimated cell type composition) that were explained by the top ten inferred PEER factors.

### *Cis*-heritability estimation of gene expression

To understand the overall contribution of *cis*-genetic variants to variation in gene expression, we only considered SNPs within 1 Mb up- and down-stream of TSS. We then employed a linear mixed model to estimate the *cis*-heritability (*cis*-*h*^2^) of each gene, while accounting for all the estimated covariates (i.e., genotype PCs and PEER factors). We estimated the genetic parameters using the restricted maximum likelihood (REML) method implemented in GCTA (v1.93.0)^88^, and defined *cis*-*h*^2^ with *P*-value < 0.05 (from the likelihood ratio) as significant. To further study the pairwise tissue similarity on gene regulation, we calculated the pairwise Pearson’s correlation of *cis*-*h*^2^ between tissues and then clustered tissues using the *hclust* function in R (v4.0.2).

### *cis*-QTL mapping for PCG, exon, lncRNA and enhancer expression

For *cis*-eQTL mapping, we followed the same pipeline as in human GTEx^7^. We first normalized the expression of PCGs across samples within each tissue using the Trimmed Mean of M-value (TMM) method, implemented in edgeR^89^, followed by inverse normal transformation of the TMM. We then performed *cis*-eQTL mapping using a linear regression model, implemented in TensorQTL (v1.0.3)^19^, while accounting for the estimated covariates (i.e., genotype PCs and the top ten PEER factors for the corresponding tissue). Within each tissue, we filtered out genes with TPM < 0.1 and/or raw read counts < 6 in more than 20% of samples. We defined the *cis*-window of PCG as ±1Mb of TSS and obtained the nominal *P*-values of *cis*-eQTL with the parameter --mode *cis_nominal* in TensorQTL. We then applied the permutation mode to calculate empirical *P*-values with parameter --mode *cis* and used the FDR method to correct the *beta* distribution-extrapolated empirical *P*-values for multiple testing. We considered genes with at least one significant (FDR ≤ 0.05) *cis*-eQTL as eGenes and genes without significant *cis*-eQTL as non-eGenes. To identify significant *cis*-eQTL associated with eGenes, we defined the empirical *P*-value of the gene that was closest to an FDR of 0.05 as the genome-wide empirical *P*-value threshold (*pt*). We obtained the gene-level threshold for each gene from the *beta* distribution by *qbeta*(*pt*, *beta_shape1*, *beta_shape2*) in R (v4.0.2), where *beta_shape1* and *beta_shape2* were derived using TensorQTL. We then considered SNPs with a nominal *P*-value below the gene-level threshold as significant *cis*-eQTL for a given gene-tissue pair.

To evaluate whether the complex genetic relatedness among samples affected *cis*-eQTL mapping using TensorQTL, we also analyzed each tissue’s data with a linear mixed model (LMM), implemented in fastGWA (v1.93.0)^90^, with a sparse genetic relationship matrix (GRM) that was constructed based on imputed genotypes for each tissue’s samples. In addition to the GRM, we also included the top ten PEER factors as above to account for the technical confounders. As done in the TensorQTL analysis, we only considered SNPs within the *cis*-window of a gene for *cis*-eQTL mapping based on fastGWA. We calculated the Pearson’s correlation of the summary statistics from fastGWA and those from the TensorQTL, including effect sizes (i.e., slope) and significance levels (i.e., −log_10_(*P*)).

Similarly, we normalized expression of exons, lncRNAs and enhancers to inverse normal transformed TMM across samples and excluded lowly expressed elements using the same approach as described above for *cis*-eQTL mapping of PCG. We then conducted *cis*-QTL mapping for exons (*cis*-eeQTL), lncRNAs (*cis*-lncQTL), and enhancers (*cis*-enQTL) using TensorQTL. For *cis*-eeQTL mapping, we defined the *cis*-window of an exon as the ±1Mb region of its source gene’s TSS. For exons, lncRNA, and enhancer *cis*-QTL mapping, we defined the *cis*-window as the ±1Mb region of the TSS of the source gene, of its TSS, and its TSS, respectively. We declared significant *cis*-QTL for exons, lncRNAs, and enhancers using the same approach as done for the *cis*-eQTL mapping of PCG. We defined exons, lncRNAs, and enhancers with at least one significant *cis*-QTL as eExon, eLncRNA, and eEnhancer, respectively.

### *cis*-sQTL mapping

We performed *cis*-sQTL mapping for genes with splicing quantifications (PSI values) and tested SNPs within ±1Mb of TSS using TensorQTL (v1.0.3)^19^, while accounting for the estimated covariates. To compute the empirical *P*-value of *cis*-sQTL, we grouped all intron clusters of a gene with the parameter: --phenotype_groups option, in the *permutation* mode of TensorQTL (v1.0.3)^19^. We then defined sGene and significant *cis*-sQTL using the same approach as used for *cis*-eQTL mapping. In addition, we defined the top *cis*-sQTL of each gene as the SNP with the lowest nominal *P*-value among all SNPs in the intron clusters for the given gene. We refer to the eGene, eExon, eLncRNA, and eEnhancer above as well as sGene collectively as eMolecule.

### Estimation of effect sizes of *cis*-eQTL

To quantitatively interpret the cellular regulatory events from the population data^91^, we calculated the allelic fold change (aFC) of a *cis*-eQTL to quantify its effect size on gene expression using aFC (v0.3)^91^, while taking account of the same covariates as done in the *cis*-eQTL mapping. We obtained the 95% confidence interval of aFC using the bootstrap method with the argument: --boot 100, and only kept *cis*-eQTL with a 95% confidence interval of aFC not overlapping with zero for further analysis.

### Conditionally independent molQTL mapping

To identify the multiple independent *cis*-QTL signals of a given molecular phenotype, we applied a forward-backward stepwise regression approach^7^, using TensorQTL (v1.0.3) with the parameter: -- mode *cis_independent* ^19^. We set the gene-level significance threshold to be the maximum *beta*-adjusted *P*-value for eGenes, sGenes, eLncRNAs, eExons and eEnhancers within each tissue after correcting for multiple testing as described above. At each iteration, we scanned the new *cis*-QTL after adjusting for all previously discovered *cis*-QTL and covariates. In addition, we employed DAP-G (v1.0.0)^92^, a Bayesian multi-SNP genetic association analysis algorithm, to fine-map the potential causal *cis*-eQTL for each eGene. The input data for the DAP-G analysis included z-scores of *cis*-eQTL and the LD matrix (genotype correlation) of genotypes in a tissue. We identified SNP clusters with independent signals, and then calculated the causal probability for each SNP cluster by summing the probabilities of individual SNPs within a cluster. We then defined SNP clusters with a causal probability of ≥ 0.8 as significant clusters.

### Internal validation of *cis*-eQTL

We conducted an internal validation of *cis*-eQTL for tissues that had sample size ≥ 80. For each tissue, we randomly and evenly divided the samples into two groups, and then conducted *cis*-eQTL mapping within each group separately, using TensorQTL. To measure the validation rate of *cis*-eQTL between groups, we calculated the π_1_ statistic, defined as the proportion of *cis*-eQTL in group *i* that are significant in group *j* ^93^, using the *qvalue* method^94^, and as well as the Pearson’s correlation of estimated effect sizes (i.e., absolute z-scores from TensorQTL) of the *cis*-eQTL between groups.

### External validation of *cis*-eQTL

To further validate the identified *cis*-eQTL, we examined two external datasets: 1) 179 animals with both 50K SNP array and blood RNA-Seq data at two time points in a composite population of ¼ Duroc, ¼ Yorkshire and ½ Landrace; 2) 100 Duroc pigs with both WGS and RNA-Seq data from muscle, liver, and duodenum.

For the first validation dataset, we analyzed all the RNA-Seq data and imputed RNA-Seq genotypes to whole-genome sequence level based on the PGRP, using the same pipeline as described above. We also imputed genotypes from the 50K SNP array to sequence level. We then conducted the *cis*-eQTL mapping at each of two time points separately based on genotypes imputed from RNA-Seq and those imputed from 50K SNP array using the same pipeline as above, while considering the first five genotype PCs and ten PEER factors as covariates. To quantify the validation rate between the validation and discovery dataset, we calculated the π_1_ statistic for the top *cis*-eQTL of genes identified in the validation dataset with those identified for blood in the PigGTEx discovery population. To further quantify the correlation of effect sizes of *cis*-eQTL between the discovery and validation datasets, we performed a meta-analysis of *cis*-eQTL using a multivariate adaptive shrinkage (MashR) method^95^. Following the same pipeline as described for the human GTEx^7^, we used z-scores (i.e., slope/slope_se from TensorQTL) of the top *cis*-eQTL of genes as input and fitted the *mash* model using 1,000,000 random SNP-gene pairs being tested as null input. We obtained the estimated effect size (i.e., the posterior mean) of the *cis*-eQTL from the *mash* function and computed Spearman’s correlation of those estimates between the discovery and validation datasets.

For the second validation dataset, we called genotypes directly from the WGS data but only considered 1,678,951 SNPs that overlapping those of the PigGTEx discovery dataset for *cis*-eQTL mapping. Using the same approach as described above, we performed *cis*-eQTL mapping for each of these three tissues using TensorQTL, while considering the first five genotype PCs and ten PEER factors as covariates. We then computed the π_1_ statistic to quantify the validation rates of *cis*-eQTL between the discovery and validation datasets.

### ASE validation of *cis*-eQTL

ASE is a complementary approach for *cis*-eQTL mapping at the individual level that is not affected by confounders among samples^96,97^. We utilized phASER (v1.1.1) to perform the haplotype-based ASE analysis^98^. First, we phased the variants from the bam and VCF files using phASER with parameters: --paired_end 1 --mapq 255 --baseq 10. We computed the mappability of each locus in the reference genome using GenMap (v1.3.0)^99^ with 75 bp k-mers, allowing two mismatches (-K 75 -E 2). We then calculated the expression of the haplotype using phASER Gene AE (v1.2.0) with default settings and generated a haplotype expression matrix across all samples using the *phaser_expr_matrix.py* script. We then used the ASE results to validate *cis*-eQTL that had ≥ 10 individuals with ASE data and ≥ 8 reads for a given gene. We first calculated the ASE-level effect size (ASE aFC) using the script *phaser_cis_var.py* with default parameters and applied 10,000 bootstraps (--bs 10000) to generate a 95% confidence interval of ASE aFC. We then calculated Spearman’s correlation between the aFC from *cis*-eQTL and from ASE at the matched loci across all 34 tissues.

### Breed-sharing patterns of *cis*-eQTL

To explore how *cis*-eQTL are shared across breeds, we divided muscle samples into eight breed groups (i.e., Asian pigs, Duroc, Yorkshire, Landrace, Landrace×Yorkshire, Duroc×Luchuan, Duroc×Pietrain, and other crossbreeds), and then performed the *cis*-eQTL mapping as described above within each breed group separately using TensorQTL (v1.0.3)^19^. We calculated the π_1_ statistic to measure the replication rate of *cis*-eQTL between breeds. In addition, to further explore the sharing pattern of *cis*-eQTL across breeds, we performed a meta-analysis of *cis*-eQTL across eight breed groups using MashR (v0.2-6)^95^ and METASOFT (v2.0.1)^20^. We used the z-scores calculated from TensorQTL (i.e., slope/slope_se) of *cis*-eQTL of genes across all 34 tissues and 8 breed groups as input for MashR (v0.2-6). To fit the *mash* model, we first randomly selected 1,000,000 SNP-gene pairs that were tested in all breed groups as null inputs. We set missing z-scores as zero with a standard error of 10^6^ *via* the *zero_Bhat_Shat_reset* = 10e6. We then obtained the estimate of effect size (i.e., the posterior mean) and the corresponding significance level (i.e., the local false sign rate, LFSR) from the *mash* function. To identify whether a *cis*-eQTL is active in a breed, we defined LFSR < 0.05 as the significance threshold, unless noted otherwise. To quantify the similarity of *cis*-eQTL effect sizes between two breeds, we calculated the Spearman’s correlation of effect size estimates of *cis*-eQTL significant (LFSR < 0.05) in at least one breed. Furthermore, we employed METASOFT (v2.0.1) to perform a meta-analysis of *cis*-eQTL across different breeds using summary statistics obtained from TensorQTL (v1.0.3) (i.e., slope and slope_se). We obtained estimates of meta-analytic effect sizes from a fixed effects model and M-values from a Markov Chain Monte Carlo (MCMC) method. The M-value represents the posterior probability that a *cis*-eQTL effect exists in a breed.

### Breed and cell type interaction *cis*-eQTL

To detect *cis*-eQTL that may be specific to a particular breed, we performed the breed-interaction *cis*-eQTL (bieQTL) analysis. We first used imputed genotypes to estimate the ancestry composition of all RNA-Seq samples across tissues using ADMIXTURE (v1.3.0) with K = 5, representing Duroc, Landrace, Yorkshire, Asian pigs, and American pigs. We only considered 33 tissues that had breed ancestry with a median ancestry proportion > 0.1 for bieQTL mapping. We performed the bieQTL mapping using the following linear regression model with an interaction term between genotype and ancestry proportion, implemented in TensorQTL (v1.0.3), separately for each tissue-breed pair:

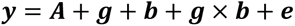

where ***y*** is the vector of gene expression values (i.e., the inverse normal transformed TMM), ***A*** represent the same covariates as in the regular *cis*-eQTL mapping, ***g*** is the genotype vector for a specific SNP, ***b*** is the proportion of a given ancestry (e.g., Duroc), ***g*** × ***b*** is the interaction term between genotype and ancestry proportion, and ***e*** is the residual error. We only considered SNPs within the *cis*-window (±1Mb of TSS) of each gene for the bieQTL mapping. We filtered out SNPs with MAF < 0.1 in the top and/or bottom 50% of samples sorted by an ancestry proportion of interest (e.g., Doruc), using TensorQTL (v1.0.3) with option: --maf_threshold_interaction 0.1. We used eigenMT^100^ in TensorQTL to correct for the multiple testing at the gene-level for the top nominal *P*- value of each gene. We then computed the genome-wide significance of genes using the Benjamini-Hochberg FDR correction on the eigenMT-corrected *P*-values. We defined genes that had at least one significant (i.e., FDR-corrected *P*-value < 0.01) bieQTL as bieGenes.

We detected cell type interaction *cis*-eQTL (cieQTL) in seven bulk tissues using the same approach as described above, where we replaced the ancestry composition with the estimated cell type composition obtained from CIBERSORTx based on single-cell RNA-Seq data from brain, lung and PBMC^74^. We considered cell types with a median enrichment percentage of > 0.1 within a tissue for cieQTL mapping. We defined genes that had at least one significant (i.e., FDR-corrected *P*-value < 0.01) cieQTL as cieGenes.

### ASE validation of ieQTL

To validate the detected ieQTL, we estimated the effect size (aFC) of the top ieQTL of ieGenes from ASE data using the script *phaser_cis_var.py* in phASER (v1.1.1)^98^. Here, we only considered ieQTL that had nominally significant ASE (*P*-value < 0.05) data in more than 10 heterozygous individuals with more than 8 reads for a gene. To filter out samples with outlier ASE values, we applied the Hampel’s test, a median absolute deviation (MAD) based method, to the allelic imbalance (AI) ratio 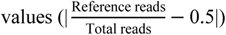 across samples^21,101^. We defined a sample as an outlier if it had |AI − median (AI)| ≥ 4.5 × MAD, where MAD = median(|AI*_i_* − median(AI)|) and AI_*i*_ is the allelic imbalance ratio value for *i*^th^ individual. We then calculated the Pearson’s correlation between aFCs of an ASE locus and ancestry/cell type proportion estimates across the remaining samples within a tissue. We considered that an ieQTL was validated by ASE data if Pearson’s correlation was nominally significant (at *P*-value < 0.05). For tissues with ≥ 5 ieQTLs, we also calculated the π_1_ statistics for significance (*P*-value) of Pearson’s correlation across breeds/cell types in a given tissue using *qvalue* method^94^ with a fixed lambda of 0.5 ^21^.

### Sharing patterns of different molQTL

To explore the specificity/similarity of the five types of molQTL, we first generated approximate LD blocks (defined by Gabriel S et al. ^102^) of SNPs across the entire genome for each tissue based on imputed genotypes using PLINK (v1.90) with parameters: --blocks no-pheno-req. For each tissue, we tested whether conditionally independent molQTL or ieQTL from different molecular phenotypes were located in the same LD block. We defined a molQTL that is unique to a molecular phenotype if it was not located in the same LD block with other types of molQTL.

### The tissue-sharing patterns of molQTL

To understand the shared or specific genetic regulatory mechanisms between tissues, we performed a meta-analysis of molQTL across all 34 tissues using MashR (v0.2-6)^95^ and METASOFT (v2.0.1)^20^ as described above. For MashR (v0.2-6), we only considered the z-scores from TensorQTL (v1.0.3) (slope/slope_se) of the top *cis*-molQTL. We obtained the estimated effect sizes (i.e., posterior means) and the corresponding significance levels (i.e., LFSR) from the *mash* function. We set LFSR < 0.05 as the significance threshold to define whether a molQTL is active in a given tissue. To estimate the pairwise tissue similarity regard to genetic regulation of gene expression, we calculated the pairwise Spearman’s correlation of effect size estimates of *cis*-molQTL between any tissue pairs, focusing on SNPs with LFSR < 0.05 in at least one tissue. For METASOFT (v2.0.1), we used summary statistics from TensorQTL (v1.0.3) (i.e., slope and slope_se) of molQTL across all tissues. We estimated the meta-analytic effect size using a fixed effect model and calculated M-values (posterior probabilities) using the MCMC method. We considered a molQTL active in a tissue when it had an M-value > 0.7. To evaluate the similarity of tissue-clustering patterns across different data types (i.e., PCG expression, splicing quantifications, exon expression, lncRNA expression, enhancer expression, *cis*-eQTL, *cis*-sQTL, *cis*-lncQTL, *cis-*eeQTL, and *cis*-enQTL), we performed k-means clustering using the *kmeans* function in the stats R package (v4.0.2), in which k (i.e., the number of clusters) was allowed to range from 2 to 20 and the maximum number of iterations was 1,000,000. We then calculated the pairwise Rand index to measure the clustering similarity using the *rand.index* function in the fossil (v0.4.0) R package (v4.0.2)^103^.

### Sequence conservation of *cis*-eQTL

To understand the evolutionarily sequence conservation of the different types of *cis*-eQTL, we downloaded PhastCons scores of 100 vertebrate species from UCSC (http://hgdownload.cse.ucsc.edu/goldenpath/hg38/phastCons100way/hg38.100way.phastCons/). We first converted the Wiggle files of PhastCons scores to bed files using the BEDOPS tool (v2.4.40)^104^, and then lifted over from human genome 38 (h38) to Sscrofa11.1 using UCSC’s LiftOver tool^105^. We used the mean phastCons scores of sequences within a gene to represent the PhastCons score of this gene. We only considered genes that had a minimum matched the length of sequences ≥ 80% of the gene’s length in LiftOver.

### Detecting *cis*-eQTL with opposite effects between tissues

To understand the tissue-specific effects of *cis*-eQTL, we examined *cis*-eQTL with opposite effects between any pair of tissues^106^. For a given tissue pair, if the top *cis*-eQTL of a given eGene were identical in the two tissues or in high LD (*r^2^* > 0.8), we designated the top *cis*-eQTL as a multi-eQTL and the eGene as a multi-eGene. We then defined the multi-eQTL whose estimated effects on the multi-eGene showed the opposite direction in the two tissues as opp-multi-eQTL and the target multi-eGene of the opp-multi-eQTL as an opp-multi-eGene.

### QTL mapping for DNA methylation (meQTL) in muscle

We conducted QTL mapping for methylation level at each CpG locus with a coverage of at least five reads in muscle, where we had 101 WGBS samples. We called SNPs from the WGBS data using *BisulfiteGenotyper* in the Bis-SNP package (v1.0.1, and dbSNP build 150) with parameters: -nt 4 - stand_call_conf 10 -mmq 30 -mbq 17 -out_modes EMIT_ALL_SITES^107^. After quality control using PLINK2^83^ (MAF ≥ 0.05, DR^2^ ≥ 0.8, and Hardy Weinberg equilibrium, HWE ≥ 1e-5), 7,410,484 SNPs remained for subsequent analysis. For methylation, we excluded CpG sites with missing rate ≥ 10% across samples. To improve the efficiency of QTL mapping, we only considered CpG loci with a deviation of methylation levels > 0.3 and a standard deviation > 0.1, yielding 18,023,521 CpG sites on 18 autosomes. We applied the rank-based inverse normal transformation to the methylation levels across samples for each CpG site and predicted 10 hidden factors using PEER (v1.3)^60^. We mapped SNPs within 1Mb around a CpG site to identify the significant SNP-CpG pairs using FastQTL (v2.184)^108^ at the threshold of *P*-value < 1e-8, considering five genotype PCs, ten PEER factors, bioproject, sex and sequencing platforms as covariates. On average, we tested 6,800 SNPs for each CpG site. We removed significant meQTL that overlapped with any CpG sites (either C or G).

### Functional enrichment analysis of molQTL

To investigate molecular mechanisms underlying the regulatory variants detected above, we examined multi-layer biological data as features, including SNPs annotated by SnpEff v.4.3^109^, sequence ontology (e.g., intron and UTR), 15 chromatin states from 14 different tissues^17^, and the DNA methylation features (i.e., HMR, ASM and meQTL) identified above. Similar to the human GTEx^7^, we employed a Bayesian hierarchical model and an expectation–maximization (EM) algorithm with the *-est* option of TORUS^110^ to explore whether the five types of identified molQTL (i.e., *cis*-eQTL, *cis*-sQTL, *cis*-lncQTL, *cis*-eeQTL, and *cis*-enQTL) were enriched for variants within these biological features. Following the human GTEx approach^7^, we examined whether the independent *cis*-eVariants were significantly enriched in the same TAD with their target eGenes. Briefly, in a tissue, we first randomly selected 1M variants from the *cis*-window of genes being tested for *cis*-eQTL as a null file. We then kept all the top variants in each *cis*-eQTL as an eQTL file. We accessed the degree of enrichment of eQTL in the same TAD with their target genes using the following formula: SameTAD ~ eQTL + |TSSdistance| + eQTL*|TSSdistance|, where SameTAD is an indicator of whether a variant-gene pair resides within the same TAD, eQTL is an indicator of whether the variant-gene pair is an eQTL or null, and |TSSdistance| is the absolute distance between the variant and the TSS of the gene. Moreover, to predict genetic variants that alter transcription factor binding sites (TFBS), we applied a custom script^111^ with TFBS models from the JASPAR (CORE 2018)^112^, HOCOMOCO (v10)^113^, and TRANSFAC (v3.2 public)^114^ databases. The TFBS models were represented as position weight matrices (PWM) and were derived from published collections of experimentally defined eukaryotic TFBS. We used the vertebrate PWM and kept only a candidate regulatory variant if the gene and the transcription factor, both impacted by the cis-eQTL, were both expressed in the same tissue.

### Integrative analysis of molQTL and GWAS of complex traits

#### GWAS summary statistics

To investigate the regulatory mechanisms underpinning complex traits in pigs, we systematically integrated the identified molQTL with summary statistics of 268 meta-GWAS from 207 complex traits of economic importance, representing five trait domains (i.e., reproduction, production, meat and carcass, health/immune, and exterior traits). In total, we performed 2,056 separate GWAS, and conducted the meta-GWAS analysis for the same traits across different populations based on GWAS summary statistics using METAL (v2011-03-25)^115^, resulting in 268 meta-GWAS results. Detailed information for each GWAS is shown in Table S17. To perform the integrative analysis of GWAS and molQTL, we overlapped significant GWAS loci with the 3,087,268 SNPs were tested in the molQTL mapping analysis, resulting in 1,507 GWAS loci with lead SNP *P*-value < 1×10^−5^.

#### Enrichment of molQTL and trait-associated variants

To examine whether molQTL were significantly enriched among the significant GWAS variants, we applied three distinct approaches as described in the following. First, we used a simple overlapping approach to examine whether a significant molQTL is more likely to be a significant trait-SNP as described in ^10^. Briefly, for each tissue, we kept SNPs with the most significant nominal *P*-value for a gene and then scaled *P*-values to a comparable level (lambda = 10) across 34 tissues. We selected the minimum *P*-value of each SNP in the 34 tissues as the background set, from which we extracted *P*-values for SNPs that overlapped with significant GWAS loci. Second, we applied QTLEnrich (v2)^7^ to quantify the enrichment degree between significant molQTL and GWAS loci. To ensure reliability of the results, we only used summary statistics of 198 GWAS for which ≥ 80% of SNPs were also tested in the molQTL mapping. Third, we applied the mediated expression score regression (MESC) method to estimate the heritability of complex trait that was mediated by the *cis-*genetic component of different molecular phenotypes (ℎ_med_^2^)^25^. We focused on the same 198 GWAS as above and calculated the LD score based on the PGRP.

#### *cis*-molQTL-GWAS colocalization

To identify shared genetic variants between the molecular phenotypes and complex traits, we performed colocalization analysis of molQTL and GWAS loci using fastENLOC (v1.0)^27^. Briefly, we obtained the probabilistic annotation of molQTL from the DAP-G (v1.0.0) and then used the *summarize_dap2enloc.pl* script to generate the annotation file of multi-tissue molQTLs. We generated approximate LD blocks across the entire genome based on the PGRP using PLINK (v1.90), as described previously^102^. We applied the TORUS tool to generate the posterior inclusion probability (PIP) of each LD block based on GWAS z-scores^110^, followed by the colocalization analysis with fastENLOC (v1.0). We obtained the regional colocalization probability (RCP) of each LD-independent genomic region using a natural Bayesian hierarchical model^116^ and considered a gene with RCP > 0.9 as significant. To identify the trait-relevant tissues, we calculated a ‘relevance score’ between a tissue and a trait by dividing the number of colocalized genes by the product of sample size and eGene proportion in this tissue. We only considered 14 tissues with a sample size greater than 100.

#### TWAS of complex traits

To explore whether the overall *cis*-genetic component of a molecular phenotype is associated with complex traits, we conducted single- and multi-tissue TWAS using S-PrediXcan^28^ and S-MultiXcan in MetaXcan (v0.6.11)^29^, respectively, based on the summary statistics of the meta-GWAS. Briefly, we employed the Nested Cross validated Elastic Net model implemented in S-PrediXcan to predict the molecular phenotype for PCG, splicing, exon, and lncRNA in all 34 tissues. To train the predictive model, we used the confounder-corrected expression or PSI values as phenotypes, and SNPs within the *cis*-windows of genes as genotypes. We kept only predictive models with cross-validated correlation ρ > 0.1 and prediction performance *P* < 0.05 for further TWAS analysis. We ran S-PrediXcan on all 268 GWAS to obtain gene-trait associations at a single-tissue level. Based on results from S-PrediXcan, we then ran S-MultiXcan to integrate predictions from multiple tissues, yielding the multiple-tissue TWAS results. We applied the Bonferroni method to correct for multiple testing and considered a corrected-*P* < 0.05 as significant.

#### MR analysis between molQTL and GWAS loci

We conducted MR analysis to infer the causality between molecular phenotypes and complex traits using SMR tool (v1.03), with genetic variants as instrumental variables^23^. We first converted the summary statistics of molQTL from TensorQTL (v1.0.3) to BESD format using SMR (v1.03) with the options: *--fastqtl-nominal-format --make-besd*. We only considered molecular phenotypes with at least one significant molQTL and top nominal *P*-value < 1×10^−5^ for the SMR test. To correct for multiple testing, we defined gene-trait pairs to pass the SMR test if the Benjamini Hochberg adjusted *P*_SMR_ < 0.05 and *P*_GWAS_ < 1×10^−5^. For gene-trait pairs that passed the SMR test, we then performed the heterogeneity in dependent instruments (HEIDI) test, with *P*_HEIDI_ ≥ 0.05 reflecting that we could not reject a single causal variant with effects on both molecular phenotype and complex trait. As a *cis*- regulator, lncRNA can regulate the expression of neighboring PCGs, and then influence complex traits. To understand this aetiology of complex traits, we performed an integrative SMR analysis that used three layers of summary-level information from *cis*-lncQTL, *cis*-eQTL and trait GWAS. We used the summary statistics of *cis*-lncQTL and *cis*-eQTL as the exposure and the outcome input for SMR (v1.03)^117^, respectively, which detected pleiotropic effects between lncRNA and PCG expression. To correct for multiple testing, we used Bonferroni correction within each tissue and defined a corrected-*P* < 0.05 as significant.

#### Comparative transcriptome between pigs and humans

We compared gene expression and its genetic regulation between pigs and humans in 17 common tissues, including adipose, artery, blood, colon, frontal cortex, heart, hypothalamus, ileum, kidney, liver, lung, muscle, ovary, pituitary, spleen, testis, and uterus. We downloaded gene expression and *cis*-eQTL data of 15,044 samples from the Human GTEx web portal (v8, https://gtexportal.org/home/) and considered genes with TPM > 0.1 as expressed. We divided genes into the following three groups based on the orthologous annotation from Ensembl (v100): one-to-one orthologous genes (n= 15,944), complex orthologous genes (n = 2,395 in pigs, and n = 2,605 in humans, 1-to-many, many-to-1, and many-to-many), and non-orthologous genes (n = 13,569 in pigs, and n = 37,651 in humans). We integrated the gene expression matrices from pigs and humans (5,173 and 15,044 samples in pigs and humans, respectively) using Seurat (v3.0), while correcting for unknown batch effects. We then visualized the divergence in gene expression between pig and human samples using *t*-SNE^49^. We detected tissue-specific genes for humans using the same method as described above for pigs. Within each tissue, we divided genes into four groups by comparing eGenes between pigs and humans: 1) eGenes shared by both species (Both), 2) eGenes only identified for humans (Human-specific), 3) eGenes only identified for pigs (Pig-specific), 4) non-eGenes in either species (Neither). To test the conservation of eGenes between pigs and humans, we did the Fisher exact test and obtained odd ratio (OR) values. To investigate the difference of these four gene groups in sequence conservation between pigs and humans, we used PhastCons scores of 100 vertebrate genomes, as described above. To explore their difference in the tolerance to loss of function mutations (LOF), we obtained the LOEUF scores from gnomAD (v2.1.1)^118^. We obtained orthologous variants between pigs and humans using LiftOver^105^, among which 112 SNPs were *cis*-eQTL in at least one tissue in humans.

#### Comparative TWAS between pigs and humans

To explore the genetic similarity of complex traits between pigs and humans, we performed a comparative analysis of TWAS summary statistics. We downloaded public human GWAS summary statistics for 136 complex traits, representing 18 trait domains (Table S27). Based on the predictive models in human GTEx v8^119^, we applied the S-PrediXcan to conduct TWAS for all 136 complex traits across 49 human tissues. We only kept TWAS results from 11 major tissues in humans that had matched tissues with sample size ≥ 100 in pigs. We only considered 15,944 1-to-1 orthologous genes. For a trait pair, we calculated the Pearson’s correlation of absolute effect size estimated of orthologous genes between pigs and humans within the matching tissue. Within each matching tissue, we used the Bonferroni correction for multiple testing and defined a corrected-*P* < 0.05 as significant.

#### Statistics & Reproducibility

No statistical method was used to predetermine the sample size. The details of data exclusions for each specific analysis are available in the Methods section. For all the boxplots, the horizontal lines inside the boxes show the medians. Box bounds show the lower quartile (Q1, the 25^th^ percentile) and the upper quartile (Q3, the 75^th^ percentile). Whiskers are minima (Q1 − 1.5 × IQR) and maxima (Q3 + 1.5 × IQR), where IQR is the interquartile range (Q3-Q1). Outliers are shown in the boxplots unless otherwise stated. The experiments were not randomized, as all the datasets are publicly available and from observational studies. The Investigators were not blinded to allocation during experiments and outcome assessment, as the data are not from controlled randomized studies.

### Data Availability

All raw data analyzed in this study are publicly available for download without restrictions from SRA (https://www.ncbi.nlm.nih.gov/sra/) and BIGD (https://bigd.big.ac.cn/bioproject/) databases. Details of RNA-Seq, WGS, WGBS, single-cell RNA-Seq and Hi-C datasets can be found in Supplementary Table 1, 2, 4, 5 and 6, respectively. All processed data and the full summary statistics of molQTL mapping are available at http://piggtex.farmgtex.org/).

### Code Availability

All the computational scripts and codes for RNA-Seq, WGS, WGBS, single-cell RNA-Seq and Hi-C datasets analyses, as well as the respective quality control, molecular phenotype normalization, genotype imputation, molQTL mapping, functional enrichment, colocalization, SMR and TWAS are available at the FarmGTEx GitHub website (https://github.com/FarmGTEx/PigGTEx-Pipeline-v0).

